# Mutational consequences of perturbing DNA repair and chromatin state in *Arabidopsis*

**DOI:** 10.64898/2026.08.04.742834

**Authors:** C. A. Meyer, R. J Schmitz

**Affiliations:** Department of Genetics, University of Georgia, Athens, Georgia, 30602

## Abstract

Mutation rate varies throughout an organism’s genome and correlates with many factors, including primary sequence, DNA methylation, gene content, chromatin accessibility, replication timing, and more. Prior work has shown that DNA repair pathways contribute to this variation by repairing certain regions more efficiently than others, sometimes through interactions with gene-associated histone modifications. However, little is known about the relative importance of different DNA repair pathways in shaping intragenomic variability in mutation rate, nor the mutational consequences of perturbing chromatin state. Here, we quantify somatic mutation rate in several DNA repair and chromatin-related *Arabidopsis* mutants using nanorate sequencing. We find that NER and MMR prevent a smaller fraction of mutations in transposable elements (TEs) compared to other regions of the genome, indicating these pathways are less efficient in heterochromatin. MMR appears to be more efficient in accessible chromatin regions, as its loss nearly abolishes the reduced mutation rate there. TC-NER is the only pathway with greater efficiency in genes than in non-genic non-TE regions, suggesting only TC-NER specifically targets genes. We assay six mutants for histone modifications/variants, but only one (*h2a.w.7*) displays an altered mutation rate. Instead, mutation rate is elevated in the chromatin remodeler mutant *ddm1* and two RNA-directed DNA methylation (RdDM) mutants. The RdDM mutants have a doubled overall mutation rate, but this increase is not localized to RdDM target regions, implicating a transcriptional change, genomic instability, or a secondary function in DNA repair.

## Introduction

Mutation rate varies not only between organisms, but also within an organism’s genome. The mutation rate of a genomic region correlates with many factors, including primary sequence (Hodgkinson and Eyre-Walker 2011; Aggarwala and Voight 2016; Supek and Lehner 2019), DNA methylation (Holliday and Grigg 1993; Weng et al. 2019), replication timing (Woo and Li 2012; Liu et al. 2013), chromatin accessibility (Polak et al. 2015; Meyer et al. 2025), gene content (Supek and Lehner 2015; Belfield et al. 2018; Gonzalez-Perez et al. 2019), and more (Gonzalez-Perez et al. 2019; Supek and Lehner 2019). Mutations arise when DNA damage is misrepaired or remains unrepaired long enough to be used as a template for DNA replication (Chatterjee and Walker 2017). Thus, intragenomic variability in mutation rate is a product of differences in DNA damage and DNA repair rates. For instance, mutation rate is higher at methylated cytosines because they are deaminated ∼2-5x faster than unmethylated cytosines (higher rate of DNA damage) and their deamination product is repaired less efficiently (lower rate of DNA repair) (Ehrlich et al. 1986; Shen et al. 1994; Lutsenko and Bhagwat 1999).

DNA repair efficiency may also explain why mutation rate correlates with replication timing and chromatin accessibility in human tumors. Early replicating regions generally have lower mutation rates than late replicating regions, but this is no longer the case in tumors with inactivated mismatch repair (MMR) (Supek and Lehner 2015). This suggests MMR is more efficient in the early stages of replication (Supek and Lehner 2015). Similarly, accessible chromatin regions are normally depleted for mutations, but this depletion is mostly lost in tumors with impaired nucleotide excision repair (NER), suggesting NER is more efficient in accessible chromatin regions (Polak et al. 2014).

Studies in both plants and animals have observed that mutation rates are lower within genes than in other regions of the genome (Supek and Lehner 2015; Krasovec et al. 2017; Belfield et al. 2018; Belfield et al. 2021; Moore et al. 2021; Monroe et al. 2022; Goel et al. 2024; Meyer et al. 2025), and this too has been linked to DNA repair. For one, the ubiquitous transcription-coupled NER (TC-NER) subpathway functions only where transcription occurs, as it detects DNA lesions which stall RNA polymerase (Duan et al. 2021). In plants, MMR is also reportedly more efficient within genes, as MMR mutants are no longer depleted for mutations in genes (Belfield et al. 2018; Quiroz et al. 2024). Lastly, human double strand break repair has been shown to differ between transcriptionally active and inactive regions, with error-free homologous recombination repair preferred over non-homologous end joining in transcriptionally active regions (Aymard et al. 2014). Intriguingly, this preference is dependent on the binding of homologous recombination repair factors to the transcription-associated histone modification H3K36me3 (Daugaard et al. 2012; Aymard et al. 2014). This suggests that histone modifications can influence mutation rate through the recruitment of DNA repair factors.

Enrichment of various histone modifications and histone variants has been found to correlate with mutation rate (Schuster-Böckler and Lehner 2012; Polak et al. 2015; Frigola et al. 2017; Monroe et al. 2022; Villalba de la Peña et al. 2023), but perturbation of these factors is necessary to establish causation. For instance, perturbation of H3K36me3 in human cells revealed that this modification also promotes MMR through an interaction with the mismatch repair protein MSH6 (Huang et al. 2018). In yeast, H3K36me3 was found to improve TC-NER efficiency (Selvam et al. 2022) and the other NER subpathway—global genome NER (GG-NER)—is dependent on H3K79me and histone acetylation (Teng et al. 2002; Tatum and Li 2011; Hodges et al. 2019). Lastly, binding assays and protein modeling of *Arabidopsis* MSH6 suggest it preferentially binds to H3K4me1 instead of H3K36me3 (Quiroz et al. 2024).

These studies show much of the intragenomic variability in mutation rate can be explained by differences in DNA repair activity across the genome, and in some cases, histone modifications serve to direct DNA repair. However, little is known about the relative importance of different DNA repair pathways in shaping intragenomic mutation rate variability, nor the mutational consequences of perturbing chromatin state.

Here, we use nanorate sequencing (NanoSeq) to identify somatic mutations in 10 repair and 11 chromatin-related *Arabidopsis* mutants. We find that NER and MMR prevent a smaller fraction of mutations in transposable elements (TEs) compared to other regions of the genome, indicating these pathways are less efficient in heterochromatin. MMR appears to be more efficient in accessible chromatin regions, as its loss nearly abolishes the reduced mutation rate there. TC-NER is the only pathway with greater efficiency in genes than in non-genic non-TE regions, suggesting only TC-NER specifically targets genes. We assay six histone modification/variant mutants, but only one (*h2a.w.7*) displays an altered mutation rate. Instead, mutation rate is significantly elevated in the chromatin remodeler *ddm1* and two RNA-directed DNA methylation (RdDM) mutants.

## Results

### Somatic mutation rate and spectrum of DNA repair mutants

We selected T-DNA mutants for several DNA repair genes (**Fig. 1A, Supplemental Table S1**): *APURINIC ENDONUCLEASE* (*ARP*, base excision repair), *DNA POLYMERASE λ* (*POLλ*, multiple pathways), *MUTS HOMOLOG 2* (*MSH2*, MMR), *MUTS HOMOLOG c* (*MSHc*, MMR), *ULTRAVIOLET HYPERSENSITIVE 3* (*UVH3*, NER), *RADIATION SENSITIVE 7A* (*RAD7A*, GG-NER), *CHROMATIN REMODELING 8* (*CHR8*, TC-NER), *KU80* (non-homologous end joining), *RADIATION SENSITIVE 5A* (*RAD5A*, post-replication repair), and *POLY-ADP-RIBOSE POLYMERASE 2* (*PARP2*, multiple pathways). NanoSeq libraries were constructed from whole shoots at flowering stage for each mutant, and a median of 633 somatic mutations were identified per genotype. As a wild-type control, we used eight previously generated (Meyer et al. 2025) and three new Col-0 libraries. The somatic mutation rate of each genotype was calculated as the number of mutations divided by the number of filter-passing sequenced bases (Methods). As expected, most mutants possessed a significantly higher somatic mutation rate than the wild type (WT), with the highest being *msh2* at 13x that of WT (**Fig. 1B**).

**Figure 1.**
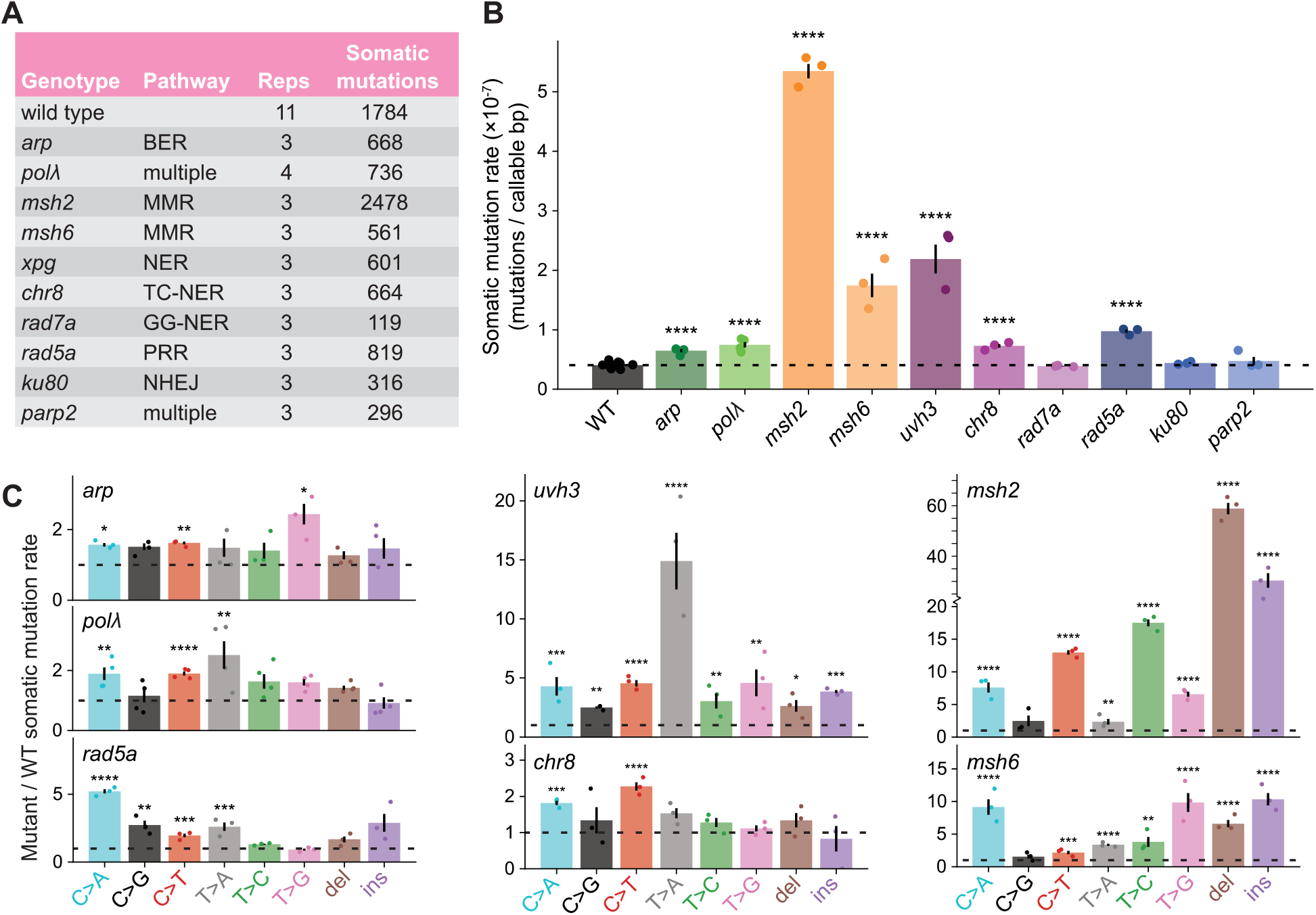
Somatic mutation rate and spectrum of DNA repair mutants. **A**, Table of DNA repair mutants sequenced with NanoSeq. BER=base excision repair, MMR=mismatch repair, TC/GG-NER=transcription-coupled/global genome nucleotide excision repair, PRR=post-replication repair, NHEJ=non-homologous end joining (see **Supplemental Table S1** for details). **B**, Somatic mutation rates of DNA repair mutants measured with NanoSeq. Somatic mutation rate was calculated as the number of mutations per filter-passing base pair sequenced. The dotted line indicates the WT somatic mutation rate. **C**, Mutation spectra of DNA repair mutants relative to WT. The somatic mutation rate of each SNV and indel type was divided by its rate in WT. The dotted line indicates a value of 1 (WT mutation rate). Only mutants with an elevated overall mutation rate are shown in C. See Supplemental Figure S3 and Supplemental Table S5 for spectra of all genotypes. **BsC**, Dots represent individual plants. Significance labels indicate the value differs from WT (Holm-adjusted t-test). *=p<0.05, **=p<0.01, ***=p<0.001, ****=p<0.0001. Error bars=±1 SEM.

Three mutants did not have an elevated mutation rate—*rad7a*, *ku80*, and *parp2*. For *rad7a*, this is likely due to the presence of two partially redundant paralogs, *RAD7B* and *RAD7C* (Lahari et al. 2018). The same may be true for *parp2*, as *PARP1* also displays PARP activity in response to DNA damage (Gu et al. 2019). For *ku80*, it may be that impaired NHEJ primarily results in large indels and structural variants, which are difficult to detect with short reads.

We analyzed the mutation spectra (rate of C>A, C>G, etc.) of the mutants with elevated mutation rates. The base excision repair mutant *arp* had increased rates of all single nucleotide variant (SNV) types, though this was only statistically significant for C>A, C>T, and T>G (**Fig. 1C**). *polλ,* potentially involved in base excision repair and other pathways (Lee et al. 2004; Roy et al. 2011; Roldan-Arjona et al. 2019), also showed significant increases in multiple SNV types (**Fig. 1C**).

In contrast, the MMR mutants *msh2* and *msh6* had a much greater increase in some SNV types than others. *msh2* displayed very high rates of C>A, C>T, T>C, T>G, and indels (primarily 1bp in length, **Supplemental Fig. S1**) but only small increases in C>G and T>A (**Fig. 1C**). In addition, C>T mutations occurred more often at (A/G)C sites than at (C/T)C sites, reversing the pattern observed in WT (**Supplemental Fig. S2**). *msh6* had increases in C>A, T>A, and T>G SNVs comparable to that of *msh2*, but neither C>T nor T>C SNVs were nearly as prevalent as in *msh2*. This contrasts with human cell lines of *msh2* and *msh6*, which have identical mutation spectra (Zou et al. 2021). Damage recognition during MMR is carried out by one of three MutS complexes, MutSα (MSH2-MSH6), MutSβ (MSH2-MSH3), or MutSγ (MSH2-MSH7) in *Arabidopsis* (Culligan and Hays 2000). Based on the mutation spectra, prevention of C>A, T>G, and T>A mutations appears to be entirely the responsibility of MutSα, whereas prevention of C>T, T>C, and indels is carried out primarily by the other complexes.

The NER mutant *uvh3* had a 2.3x increase in T>A mutations and a more modest increase in all other SNV types, insertions, and deletions (**Fig. 1C**). The strong T>A signature occurred primarily at ATA, GTA, and TTA sites (**Supplemental Fig. S3B**), a pattern previously observed under UV treatment and attributed to thymidylyl-(3′–5′)-deoxyadenosine (TA*) lesions (Zhao and Taylor 1996; Nakamura et al. 2021; Meyer et al. 2025). *chr8* (TC-NER) had significantly elevated rates only of C>A (1.8x) and C>T (2.3x) SNVs, which were only a portion of those seen in *uvh3* (4.3x C>A and 4.6x C>T). This is consistent with TC-NER performing a subset of NER while GG-NER does the rest. The lack of a strong T>A signal in *chr8* suggests either TA* does not stall RNA polymerase and initiate TC-NER or GG-NER is more efficient at detecting TA* than other damage types.

Lastly, the post-replication repair mutant *rad5a* shows a strong C>A signature (**Fig. 1C**). This repair pathway bypasses lesions that stall the replication fork using one of two mechanisms. The first is translesion synthesis, which uses a translesion polymerase to replicate across the lesion but is error prone (Gao et al. 2017). The second is template switching, which is error free but requires an undamaged DNA strand to use as a template (Gao et al. 2017). The yeast ortholog of *RAD5A* directs repair toward the template switching mechanism (Chen et al. 2008), so we suspect mutation rate is higher in *rad5a* because of increased translesion synthesis. The C>A signature of *rad5a* does not match the error profiles of most yeast and human translesion polymerases, which show enrichment for T>C errors (Pol η, ζ, and θ) or little enrichment for any single error (Pol κ) (Ohashi et al. 2000; Zhong et al. 2006; McCulloch et al. 2007; Arana et al. 2008). However, the translesion polymerases PRIMPOL and REV1 contribute to a C>A signature in human cells (García-Medel et al. 2021; Gyüre et al. 2023). Thus, the *rad5a* mutation rate increase may arise from the activity of these two translesion polymerases during post-replication repair.

### MMR prevents more early/meristematic mutations

Somatic mutations which occur early in development and/or within the meristem will expand into large clonal sectors within the plant (Schoen and Schultz 2019). This makes early/meristematic mutations more likely to be detected in multiple NanoSeq molecules than mutations which occur in differentiated tissues (**Fig. 2A**, one molecule equals the consensus of multiple PCR duplicate reads). In all genotypes, most mutations were found in only a single molecule (**Fig. 2B,C**). As we sequenced a median of 192 molecules per site, these single-molecule mutations have an observed abundance of ∼1/192. However, we did not sequence all molecules within the plant, so the true abundance of many single-molecule mutations is likely much lower (**Fig. 2D**).

**Figure 2.**
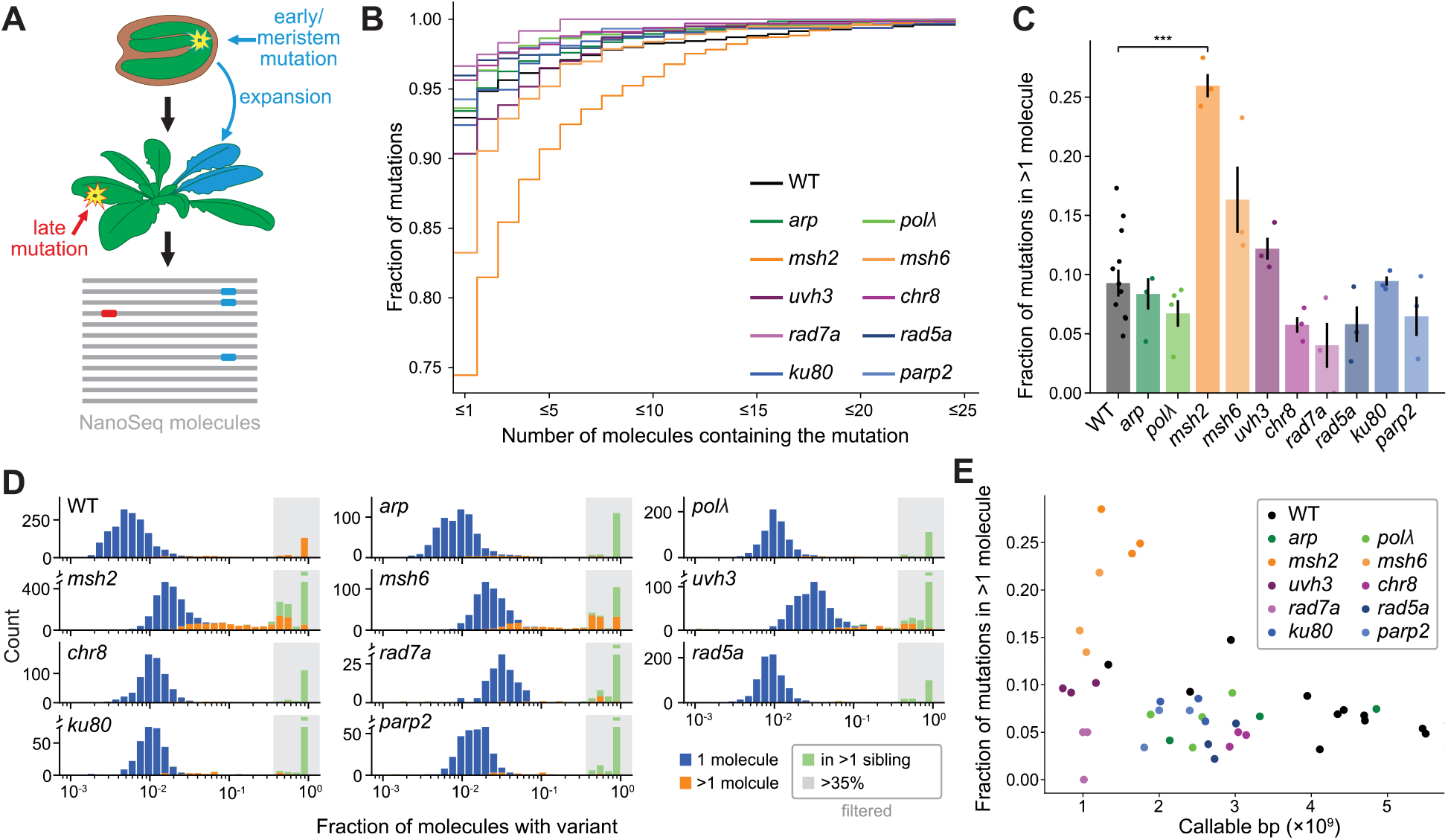
MMR prevents more high-abundance mutations. **A**, Illustration of how early (blue) and late (red) occurring mutations yield different mutation abundances in the plant. Mutations arising early in development or within meristematic tissues (top) will expand into larger clonal sectors in the adult plant (middle) and are more likely to be detected in multiple NanoSeq molecules (bottom). In contrast, mutations arising late in development or in terminal tissues will have limited to no expansion, making them unlikely to be detected in multiple molecules. **B**, Cumulative distribution of mutation abundance in the DNA repair mutants. A point on the graph indicates the fraction of mutations supported by a certain number of NanoSeq molecules or fewer. *msh2* has a greater proportion of high-abundance mutations compared to the other genotypes. **C**, Fraction of mutations supported by >1 NanoSeq molecule. Dots represent individual plants. Error bars=±1 SEM. Significance labels indicate Holm-adjusted t-tests. **D**, Mutation abundance histograms. Distribution of mutation abundance calculated as the fraction of overlapping NanoSeq molecules which support a mutation. Mutations which were found in multiple sibling plants (green) or at an abundance >35% (shaded area) were considered germline mutations and not included in any other analysis. The >1 molecule mutations in *msh2* are at frequencies far lower than expected of heterozygous variants (0.5). **E**, Scatterplot of the fraction of high abundance mutations and callable bases (the number of sequenced bases in the library passing all filters). Sequencing coverage does not explain the high rate of >1 molecule mutations in *msh2*.

We noted that a greater proportion of the mutations in *msh2* were supported by multiple DNA molecules compared to WT (p=0.00024 Holm-adjusted t-test, **Fig. 2B,C**). These high-support *msh2* mutations are not heterozygous germline mutations, as they appear at frequencies far below 0.5 (**Fig. 2D**). They also cannot be explained by sequencing depth, as the *msh2* samples were sequenced to a lower depth than most genotypes, so we would expect to detect fewer, not more, mutations in multiple molecules (**Fig. 2E**). In addition, there is little to no correlation between the proportion of high-support mutations and sequencing depth, mutation rate, or number of mutations in the non-*msh* samples (**Supplemental Fig. S4**). Thus, we conclude that loss of MMR leads to a disproportionate increase in early/meristematic mutations, while loss of other repair genes does not.

### MMR and NER prevent a greater fraction of mutations in non-TE regions

To determine whether any repair pathways were more efficient in certain regions of the genome, we divided the genome into three categories—genic, TE, and other (non-genic non-TE) regions. In the WT, mutation rate was lowest in genes, slightly higher in non-genic non-TE regions, and highest in TEs (**Fig. 3A**). This could not be explained by purifying selection, as we observed a dN/dS of 1.02 across the somatic mutation dataset (**Supplemental Fig. S5**). For all repair mutants, TEs still had the highest mutation rate (**Fig. 3A**). However, when quantifying how much mutation rate increased for each region relative to WT, we found that multiple DNA repair mutants displayed a greater increase in genic and non-genic non-TE regions than in TE regions (**Fig. 3B**). This increase was observed in *msh2*, *msh6*, and *uvh3* but was only statistically significant for *msh2* (p=0.0014, 0.32, and 0.37, Holm-adjusted t-test), suggesting MMR and potentially NER are more efficient within euchromatic regions.

**Figure 3.**
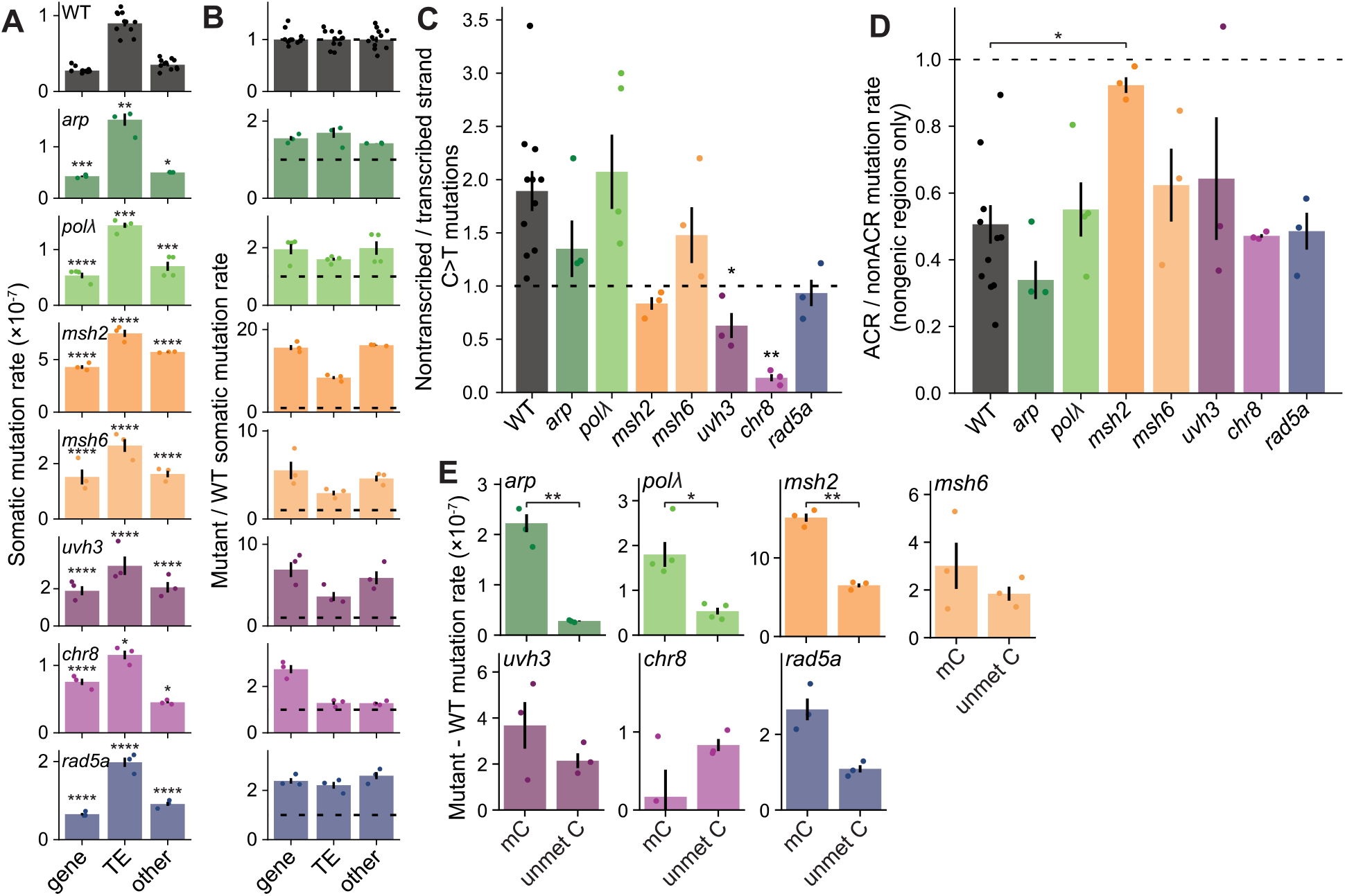
Regional biases of DNA repair mutants. **A**, Somatic mutation rate in genes, TEs, and other (non-genic and non-TE) regions. Only mutants with an elevated overall mutation rate are displayed. Significance labels indicate the value differs from WT. Mutation rate is highest in TE regions for all genotypes, and genes have a lower mutation rate than non-genic non-TE regions for all genotypes except *chr8*. **B**, Somatic mutation rate of each genotype relative to that of WT in the same region. Dotted lines represent a value of one (WT mutation rate). MMR and NER prevent a higher proportion of mutations in non-TE regions. **C**, Ratio of C>T mutations where the cytosine is on the non-transcribed strand vs on the transcribed strand of genes. Significance labels indicate the value differs from WT. The strand asymmetry flips to being more prevalent on the transcribed strand in the TC-NER mutant. **D**, Mutation rate in accessible chromatin regions (defined as WT ATAC-seq peaks) divided by non-accessible chromatin regions. Bases within genes were omitted from the analysis. Loss of MMR nearly abolishes the depletion of mutations in accessible chromatin regions. **E**, Mutation rate excess (mutant – WT somatic mutation rate) at methylated and unmethylated cytosines. Methylated cytosines were identified from WT whole genome bisulfite sequencing data. Most repair pathways have a higher rate of excess mutations at methylated cytosines. **A–E**, Dots represent individual plants. Significance labels indicate Holm-adjusted t-tests. *=p<0.05, **=p<0.01, ***=p<0.001, ****=p<0.0001. Error bars=±1 SEM.

For *chr8*, mutation rate increased primarily within genes, as expected for a TC-NER mutant. This made *chr8* the only repair mutant with a higher mutation rate in genes than in non-genic non-TE regions. There was a slight, but significant increase in the mutation rate of TEs and non-genic non-TE regions (p=0.040, 0.047, Holm-adjusted t-test), potentially reflecting small amounts of transcription—and thus TC-NER activity—outside of annotated protein-coding genes. As TC-NER recognizes lesions specifically in the transcribed strand, it produces a bias in mutation rates between the transcribed and non-transcribed strands. This can be seen in C>T and C>G mutations in WT plants, which have different frequencies depending on which strand the cytosine is located (p=0.0009 & p=0.028, Holm-adjusted t-test, **Supplemental Fig. S6**). In the WT, C>T mutations are 1.9x more frequent when the cytosine is on the non-transcribed strand. *chr8* displayed a reversal of this pattern, with cytosines on the transcribed strand now having 7.2x the rate of the other (**Fig. 3C**). This suggests either that DNA damage is more common at transcribed strand cytosines or, more plausibly, that damage on the transcribed strand is especially mutagenic when TC-NER is impaired. Three other mutants, *msh2*, *uvh3*, and *rad5a*, also had diminished or reversed strand bias in C>T mutations, though this could be explained by a higher overall mutation rate drowning out the signal of TC-NER.

Intergenic accessible chromatin regions are depleted for mutations in WT plants (Meyer et al. 2025). To test whether this is dependent on DNA repair activity, we compared mutation rates in intergenic accessible and non-accessible chromatin (**Fig. 3D**). In *msh2*, mutation rate was nearly equal between accessible and non-accessible chromatin regions, suggesting MMR alone is responsible for the reduced mutation rate in accessible chromatin regions.

Lastly, to test whether the damage repaired by each pathway is influenced by DNA methylation, we calculated the excess mutation rate (mutant minus WT mutation rate) in methylated and unmethylated cytosines for each mutant. For most mutants, excess mutation rate was higher at methylated cytosines (**Fig. 3E**), though this difference was only statistically significant for *arp* (p=0.007), *polλ* (p=0.033), and *msh2* (p=0.002, Holm-adjusted t-test). This suggests that DNA damage repaired by each of these pathways occurs more frequently or is more mutagenic at methylated cytosines.

### Somatic mutation rate of chromatin-related mutants

Work in other organisms has established links between DNA repair and histone modifications/variants (Teng et al. 2002; Tatum and Li 2011; Daugaard et al. 2012; Aymard et al. 2014; Huang et al. 2018; Hodges et al. 2019; Selvam et al. 2022; Quiroz et al. 2024). To investigate how such relationships might influence mutation rate, we performed NanoSeq on mutants for several chromatin-related genes (**Fig. 4A**): *SET DOMAIN GROUP 8 (SDG8), CURLY LEAF (clf), SU(VAR)3-9 HOMOLOG 4/5/6 (SUVH4/5/6), H2A.X.3/5, H2A.W.7, PHOTOPERIOD-INDEPENDENT FLOWERING 1 (PIE1), DNA METHYLTRANSFERASE 1 (MET1)* epiRIL, *DECREASED DNA METHYLATION 1 (DDM1), NUCLEAR RNA POLYMERASE E1* (*NRPE1)*, and *NUCLEAR RNA POLYMERASE D1 (NRPD1)*. The histone methyltransferase mutants *sdg8* and *clf* are partially depleted for H3K36me3 and H3K27me3 (Schubert et al. 2006; Jiang et al. 2008; Xu et al. 2008; Shu et al. 2019; Zhao et al. 2019), while *suvh4/5/6* loses all or nearly all H3K9me2 (Ebbs and Bender 2006; Tsuchiya and Eulgem 2013; Yu et al. 2017). The histone variant mutants *H2A.X.3/5* and *H2A.W.7* are direct knockouts of histone genes (i.e., complete loss of H2A.X and H2A.W.7), whereas the chromatin remodeler mutant *pie1* is only partially depleted for H2A.Z (Deal et al. 2007; Luo et al. 2020). The *met1* epiRIL is the product of a cross between WT and *met1* and has functional *MET1* but lacks mCG in large portions of the genome (Reinders et al. 2009). *DDM1* is a chromatin remodeler with roles in H2A.W deposition and maintenance of DNA methylation in heterochromatin (Zemach et al. 2013; Osakabe et al. 2021). *NRPE1* and *NRPD1* are components of RNA polymerase V and IV, which function in RNA-directed DNA methylation (Matzke and Mosher 2014).

**Figure 4.**
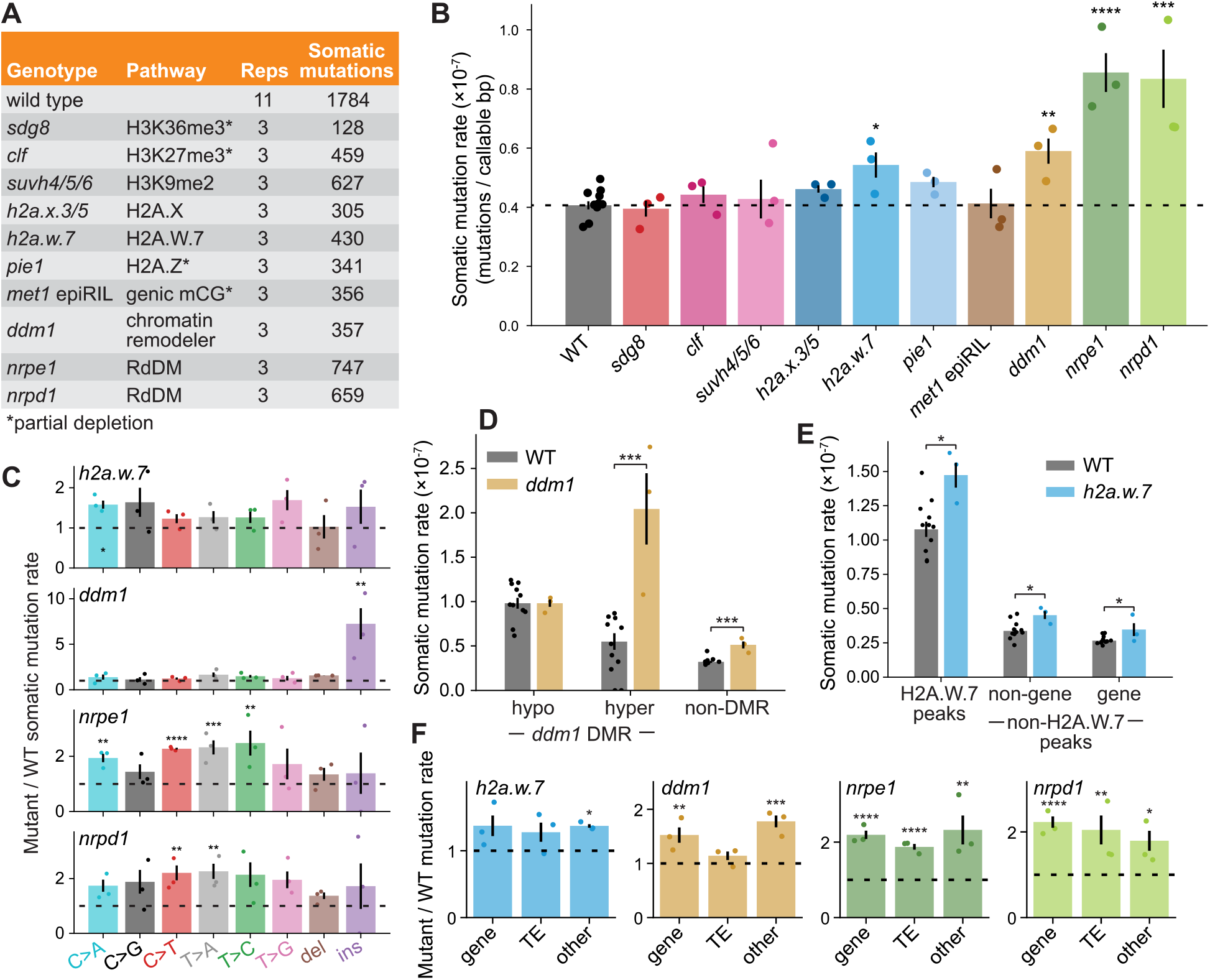
Elevated mutation rate in *h2a.w.7*, *ddm1*, and RdDM mutants. **A**, Table of chromatin-related mutants sequenced with NanoSeq. *met1* epiRIL= no CG methylation in some genomic windows, RdDM=RNA-directed DNA methylation. **B**, Somatic mutation rates of chromatin mutants measured with NanoSeq. The dotted line indicates the WT somatic mutation rate. Significance labels indicate the mutation rate differs from WT. **C**, Somatic mutation rate of each SNV and indel type divided by its rate in WT. Significance labels indicate the mutation rate of that SNV/indel differs from the WT. The dotted line indicates a value of one (WT mutation rate). *ddm1* shows a large increase in insertion rate. **D**, Somatic mutation rate in *ddm1* hypomethylated DMRs, *ddm1* hypermethylated DMRs, and non-DMR regions. Mutation rate in *ddm1* is not increased in regions where DDM1 is required for methylation maintenance. **E**, Somatic mutation rate in H2A.W.7 ChIP-seq peaks, non-genic sites outside of H2A.W.7 peaks, and genic sites outside of H2A.W.7 peaks. The mutation rate increase in *h2a.w.7* is not restricted to H2A.W.7 peaks. **F**, Somatic mutation rate in genes, TEs, and other (non-gene and non-TE) regions divided by the WT mutation rate in that region. Mutation rate is elevated primarily in non-TE regions of *ddm1* but has no bias in *h2a.w.7*, *nrpe1*, or *nrpd1*. **B-F**, Dots represent individual plants. Significance labels indicate Holm-adjusted t-tests. *=p<0.05, **=p<0.01, ***=p<0.001, ****=p<0.0001. Error bars=±1 SEM.

Six of the ten chromatin mutants had no overall increase in mutation rate (**Fig. 4B**). To test for a localized mutation rate increase in these mutants, we compared mutation rate in ChIP-seq peaks of the affected modification/variant to other regions of the genome (**Supplemental Fig. S7, Supplemental Fig. S8**). Even with this approach, we found that mutation rate was not significantly altered within H3K27me3 peaks in *clf*, H3K9me2 peaks in *suvh4/5/6*, H2A.X peaks in *h2a.x.3/5*, nor H2A.Z peaks in *pie1* (**Supplemental Fig. S7**). Despite H3K36me3 having a reported role in mammalian DNA repair (Daugaard et al. 2012; Aymard et al. 2014; Huang et al. 2018), *sdg8* also did not have an increased mutation rate in regions with diminished H3K36me3. However, *sdg8, clf, and pie1* still possess a substantial quantity of their associated marks, so it remains to be seen whether complete knockouts would have altered mutation rates.

*h2a.w.7, ddm1*, *nrpe1*, and *nrpd1* all had elevated overall mutation rates (**Fig. 4B**). To investigate why, we compared mutation spectra of these mutants to the WT (**Fig. 4C**). We noted that the rate of insertions increased far more than other mutation types in *ddm1*. Unlike in the WT, these insertions were mostly longer than 1 bp (**Supplemental Fig. S1**) and some appeared to be structural variants (**Supplemental Fig. S9**). To see if mutation rate increased specifically where *ddm1* functions in DNA methylation maintenance, we identified differentially methylated regions (DMRs) in *ddm1* compared to WT using existing bisulfite sequencing data (**Supplemental Fig. S10**) (Stroud et al. 2013). We found that *ddm1* mutation rate was greatly increased in *ddm1* hypermethylated DMRs, moderately increased in non-DMRs, and unchanged in *ddm1* hypomethylated DMRs (**Fig. 4D**). This was true even when insertions were not included in the analysis (**Supplemental Fig. S11**). Thus, it appears that DDM1 activity prevents mutations even where it is not involved in DNA methylation maintenance.

In *h2a.w.7*, *nrpe1*, and *nrpd1*, no single mutation type was obviously increased more than others. Surprisingly, the mutation rate increase in *h2a.w.*7 was not limited to H2A.W-marked regions (**Fig. 4E**), nor was it biased toward genes or TEs (**Fig. 4F**). Thus, depletion of H2A.W seems to increase mutation rate globally.

### RdDM mutants display a global increase in mutation rate

Mutation rate was increased two-fold in the RdDM mutants *nrpe1* and *nrpd1*. The RdDM pathway establishes and maintains DNA methylation at target loci using a small RNA feedback loop (Matzke and Mosher 2014). NRPD1 is a subunit of RNA PolIV, which transcribes 24 nucleotide small RNAs from target loci. These small RNAs hybridize to long noncoding transcripts produced by PolV (with subunit NRPE1) and recruit cytosine methyltransferases. We anticipated that the mutation rate increase in *nrpe1* and *nrpd1* would be localized to RdDM target regions, so we identified RdDM targets as the union of NRPE1 ChIP-seq peaks, *nrpe1* hypomethylated DMRs, and *nrpd1* hypomethylated DMRs. Unexpectedly, mutation rate increased more outside of RdDM targets than within them for both *nrpe1* and *nrpd1* (**Fig. 5A**, **Supplemental Fig. S12**).

**Figure 5.**
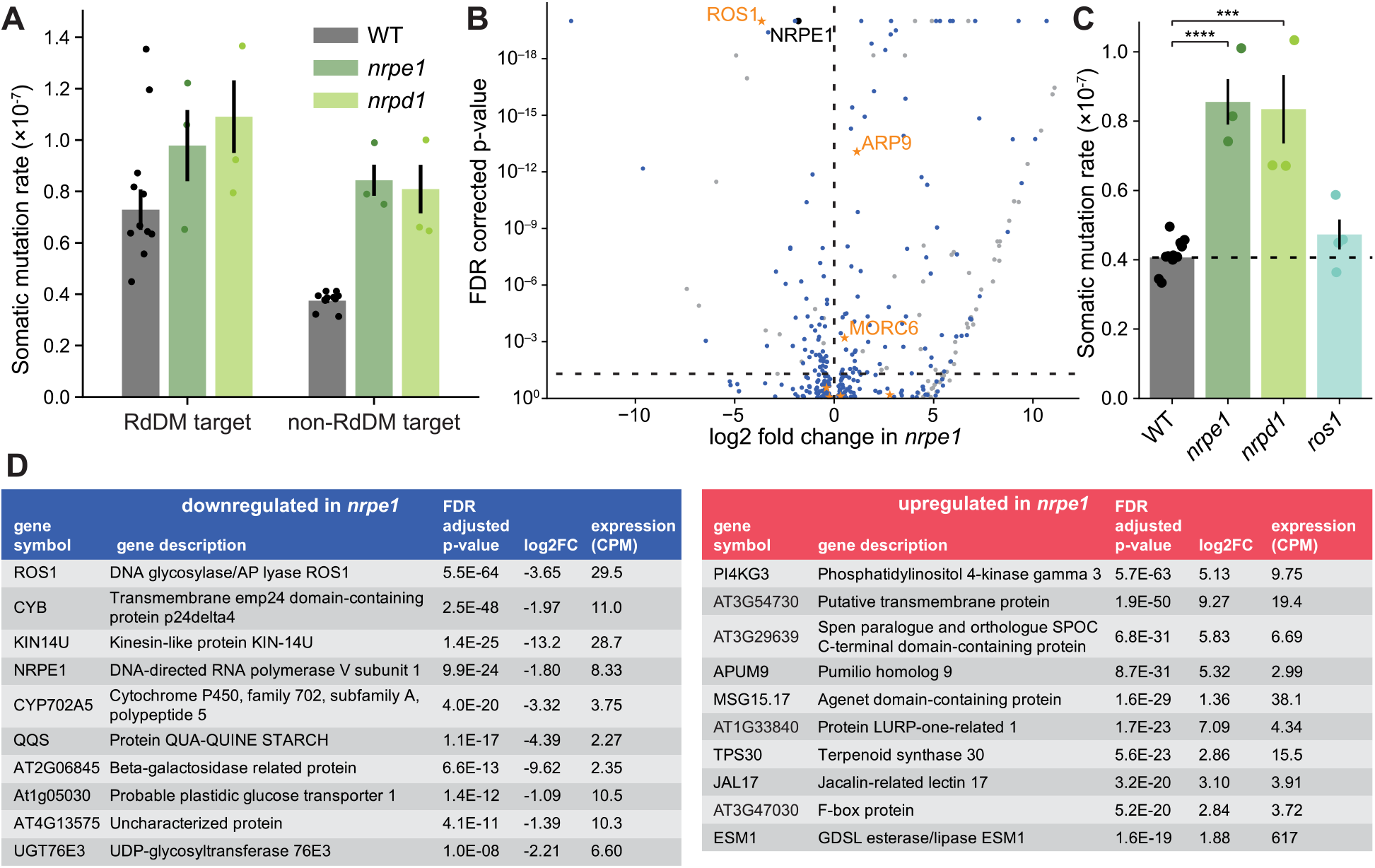
Increased mutation rate in RdDM mutants is global and not due to *ROS1* downregulation. **A**, Somatic mutation rate in RdDM targets and other sites. RdDM targets were defined as the union of NRPE1 ChIP-seq peaks, *nrpe1* hypomethylated DMRs, and *nrpd1* hypomethylated DMRs. Mutation rate is increased both within and outside of RdDM target regions in *nrpe1* and *nrpd1*. **B**, Volcano plot of differentially expressed genes in *nrpe1*. Genes upregulated in *nrpe1* are to the right of the origin. Horizontal line indicates statistical significance (FDR corrected p value < 0.05). Non-protein coding genes are colored grey and *NRPE1* is colored black. Genes annotated with a “DNA repair” or “DNA damage” GO term are represented with orange stars and labeled with the gene name. **C**, Somatic mutation rate in the catalytic mutant *ros1-7* and RdDM mutants. Significance labels indicate Holm-adjusted t-tests. *=p<0.05, **=p<0.01, ***=p<0.001, ****=p<0.0001. Mutation rate is not elevated in *ros1*. **D**, Top 10 differentially downregulated and upregulated genes in *nrpe1*. The DNA glycosylase *ROS1* is the top downregulated gene. **AsC**, Dots represent individual plants. Error bars=±1 SEM.

As this global mutation rate increase could result from a change in the expression of a DNA repair or DNA damage related factor, we analyzed existing *nrpe1* RNA-seq data. We identified 123 upregulated and 65 downregulated genes in *nrpe1* (FDR-corrected p-value < 0.05, **Fig. 5B,D**, **Supplemental Table S2**). 39 of the upregulated and 9 of the downregulated genes were annotated as transposable element genes. We searched for any differentially expressed genes annotated with a gene ontology term containing “DNA repair” or “DNA damage” and identified two upregulated (*MORCc* and *ARPS*) and one downregulated (*ROS1*) gene. No gene ontology terms were significantly enriched in the upregulated or downregulated genes (**Supplemental Table S2**).

The repair gene with the greatest expression change was *ROS1*, a DNA glycosylase with a primary function of excising 5mC but with *in vitro* activity for removing G·T mispairs (Agius et al. 2006; Ponferrada-Marín et al. 2009; Ponferrada-Marín et al. 2011). To test whether reduced *ROS1* expression could explain the elevated mutation rate in *nrpe1* and *nrpd1*, we performed NanoSeq on the catalytic missense mutant *ros1-7*. We found that mutation rate was not significantly increased in *ros1-7*, suggesting *ROS1* downregulation is not causing the elevated mutation rate in *nrpe1* and *nrpd1* (**Fig. 5C**).

## Discussion

### Repair functions of *ARP* and the MMR complexes

This study provides an opportunity to evaluate the importance of DNA repair pathways in maintaining genome integrity under standard conditions. For instance, the base excision repair protein ARP is an AP endonuclease which resolves abasic sites generated spontaneously or after excision of damaged bases by monofunctional DNA glycosylases (Roldan-Arjona et al. 2019). Mammals have only one AP endonuclease gene, and its loss is lethal (Meira et al. 2001; Fung and Demple 2005). While *Arabidopsis* has three AP endonucleases and *arp* plants are viable, they are more sensitive to multiple genotoxic agents (Córdoba-Cañero et al. 2011; Akishev et al. 2016). In addition, ARP is the only AP endonuclease with detectable *in vitro* activity on abasic site analogs, and this activity is lost in *arp* cell extracts (Córdoba-Cañero et al. 2011; Lee et al. 2014). Given that abasic sites are estimated to occur hundreds or thousands of times per day in mammalian cells (Kim and Wilson 2012), we expected *arp* plants to have a much higher mutation rate.

Instead, *arp* mutation rate was only 1.6x that of WT, suggesting that—despite the importance of *ARP* under genotoxic stress and the apparent absence of redundancy *in vitro*—abasic sites are generally properly repaired in *arp* under standard conditions. This may be because there is sufficient redundancy *in vivo* with the other AP endonucleases or other classes of endonuclease, as is seen in yeast with the NER endonuclease *RAD1* (Guillet and Boiteux 2002).

Using *msh2* and *msh6* mutants, we observed a strong partitioning of repair activity between the MutSα (MSH2-MSH6) and the MutSβ (MSH2-MSH3) and MutSγ (MSH2-MSH7) complexes. Prevention of C>A, T>A, and T>G mutations could be attributed entirely to loss of MutSα, as they were equally elevated in *msh2* as in *msh6*. Prevention of C>T, T>C, and indels was instead attributed to MutSβ and MutSγ, as they were elevated primarily in *msh2*. Thus, it appears that transition mismatches (C>T & T>C) are not repaired by MutSα. This contrasts with yeast and human, where recognition of base mismatches is performed almost exclusively by MutSα, recognition of >1 base indel loops is performed by MutSβ, and single base indel loops are recognized by both (Marti et al. 2002). As *MSH7* is a paralog of *MSH6* unique to plants (Culligan and Hays 2000), we suspect mammalian MutSα activity is partitioned between MutSα and MutSγ. *In vitro* assays of *Arabidopsis* MutSα and MutSγ binding show MutSγ has a strong preference for mispairs over C·G pairs but MutSα does not (Wu et al. 2003). In addition, ∼77% of replicative polymerase (B family) errors are C>T or T>C mismatches (Betancurt-Anzola et al. 2025), the SNV types our data suggest are repaired primarily by MutSγ/MutSβ. Thus, we propose that MutSγ is primarily responsible for repair of DNA polymerase errors, while MutSα may instead repair other lesions, such as 8-oxoGuanine (C>A), which can be repaired by mammalian and yeast MutSα (Edelbrock et al. 2013).

*msh2* uniquely displayed a higher fraction of high abundance mutations than the other genotypes. This suggests MMR prevents more mutations early in development and/or within meristematic tissues, whereas the other pathways prevent more mutations in terminal tissues. This seems to match MMR’s primary function in repairing polymerase errors made during DNA replication, which occurs in the meristem but stops in terminal tissues. The meristem may also be protected from certain types of DNA damage repaired by other pathways; oxidative damage is less common in non-photosynthesizing tissues like the meristem, and the meristem is not directly exposed to ultraviolet light (Foyer 1996; Noctor et al. 2002; Lodeyro et al. 2016). These damage sources are mostly repaired by base excision repair and NER (Barnes and Lindahl 2004; Canturk et al. 2016). The hypothesis that MMR is more active in early/meristematic tissues is further supported by existing mutation accumulation line data. Belfield et al. (2018) measured germline mutation rates of *msh2* and reported a similar mutation spectrum to our somatic mutation data. However, they observed a ∼170x mutation rate increase in *msh2*, whereas we observed only a 13x increase. Mutation accumulation lines detect mutations along the germline but not in terminal tissues, so an increased importance of MMR in germline cells might explain this discrepancy.

### MMR and NER are less efficient in heterochromatic regions

Previous studies in *Arabidopsis* and other species have noted elevated mutation rates in TEs and reduced mutation rates in genes and accessible chromatin regions (Weng et al. 2019; Monroe et al. 2022; Quiroz et al. 2024; Meyer et al. 2025). Here, we find that MMR and NER prevent a smaller fraction of mutations in TEs than in non-TE regions, suggesting they are less efficient in heterochromatin. Prior work in *Arabidopsis* has reported that MMR preferentially protects genes from mutation (Belfield et al. 2018; Quiroz et al. 2024).

However, in our MMR mutant and NER mutant, mutation rate increased by the same amount in genes as in non-genic non-TE regions. We only observed a gene-specific effect in the TC-NER mutant. Thus, it appears that TC-NER is the only assayed pathway that specifically targets genes, whereas MMR and GG-NER, the other NER subpathway, have reduced activity in heterochromatic regions.

Loss of MMR nearly equalized the mutation rate between accessible chromatin regions and non-accessible chromatin regions. No other repair mutant had this effect, contrasting with evidence that NER plays this role in human tumors (Polak et al. 2014). Repair of DNA damage is known to be impaired within nucleosome-bound DNA for MMR (Li et al. 2009; Goellner 2020), NER (Wang et al. 1991; Ura et al. 2001) and base excision repair (Beard et al. 2003; Cole et al. 2010). Thus, it is surprising that we did not see more repair pathways with increased efficiency in accessible chromatin regions.

Work in human cells has noted a role for H3K36me3 in promoting homologous recombination repair (Daugaard et al. 2012; Aymard et al. 2014) and MMR (Li et al. 2013; Huang et al. 2018) through interactions with chromatin reader domains. We found that partial depletion of H3K36me3 in *sdg8* had no effect on mutation rate in *Arabidopsis*. Quiroz et al. (2024) hypothesized that in plants, H3K4me1 takes over this role. However, recruitment to H3K4me1 could not explain our observation that MMR and NER appear just as efficient in genes as in non-genic non-TE regions (which lack H3K4me1).

Transcription is thought to increase rates of DNA damage by leaving the non-transcribed strand in a vulnerable single-stranded state, promoting R-loop formation, and causing transcription-replication conflicts (Jinks-Robertson and Bhagwat 2014). This increase in damage is offset by the activity of TC-NER within transcribed strands. We observed that in WT plants, cytosines on the non-transcribed strand were 1.9x as likely to mutate to thymine, consistent with TC-NER repairing lesions at transcribed strand cytosines—likely cyclobutane pyrimidine dimers. Loss of TC-NER reversed this effect, with transcribed strand cytosines now 7.2x more likely to mutate to thymine. We suspect this is not due to increased DNA damage on the transcribed strand, as it conflicts with our understanding of transcription-associated mutagenesis, where the non-transcribed strand is more susceptible to multiple sources of DNA damage (Francino and Ochman 2001; Klapacz and Bhagwat 2005; Fix et al. 2008; Jinks-Robertson and Bhagwat 2014). In addition, cyclobutane pyrimidine dimers form at nearly equal rates between the two strands in yeast and human cells (Mao et al. 2016; Heilbrun et al. 2021). Instead, we hypothesize that DNA damage is more mutagenic (rather than more common) in the transcribed strand when TC-NER is absent. This could occur if stalled RNA polymerase obscures the lesion from repair or increases the chances of transcription-replication conflicts.

### Mutation rate increased in *h2a.w.7*, *ddm1*, and RdDM mutants but not histone modification mutants

We found that partial depletion of H3K36me3 (*sdg8*), H3K27me3 (*clf*), or H2A.Z (*pie1*) did not alter mutation rate, nor did complete loss of H3K9me2 (*suvh4/5/6*) or H2A.X. This was especially surprising for H2A.X given its role in recruiting repair factors to double strand breaks after phosphorylation to γH2A.X (Fernandez-Capetillo et al. 2004; Donà and Mittelsten Scheid 2015). We observed a small mutation rate increase in *h2a.w.7*, which is also phosphorylated in response to double strand breaks in *Arabidopsis* (Lorković et al. 2017). However, the increased mutation rate wasn’t localized to H2A.W.7 peaks. One possible explanation for this is redundancy between H2A.X and H2A.W.7. In complete H2A.W knockouts (*h2a.w.6/7/12*), H2A.X replaces H2A.W in WT H2A.W peaks (Bourguet et al. 2021). In addition, the dephosphorylation rate of H2A.X and H2A.W.7 is dependent on whether the other variant is missing (Lorković et al. 2017). If H2A.X and H2A.W.7 act redundantly, it may explain why mutation rate was not increased in *h2a.x.3/5*, as its function is covered by H2A.W.7. The global mutation rate increase in *h2a.w.7* could be due to dilution of H2A.X in its normal regions.

Mutation rate was elevated in *ddm1*, which saw high rates of insertions and likely structural variants. DDM1 deposits H2A.W (Osakabe et al. 2021), but since *h2a.w.7* did not share the extra insertion phenotype, we suspect another function explains the mutation rate. LSH/HELLS (the human ortholog of DDM1) has known roles in double and single strand break repair (Burrage et al. 2012; Peixoto et al. 2022; Joseph et al. 2025). The *Neurospora crassa* DDM1 homolog appears to instead function in stabilization of replication forks when repair intermediates are encountered (Basenko et al. 2016). *Arabidopsis ddm1* shows sensitivity to the DNA damaging agent methylmethane sulphonate and increased accumulation of UV-B damage (Yao et al. 2012; Qüesta et al. 2013), both of which can stall replication forks (Rupp and Howard-flanders 1968; Groth et al. 2010). Our observation of many insertions and structural variants in *ddm1* is consistent with a role of DDM1 in double strand break repair or stabilization of replication forks, and this activity seems to be important genome-wide rather than exclusively within heterochromatin.

Lastly, mutation rate was roughly doubled in the RdDM mutants *nrpd1* and *nrpe1*. This increase was not restricted to RdDM targets, making us suspect an expression change of a DNA repair or DNA damage factor. However, mutants of the most promising downregulated candidate, *ROS1,* did not display an elevated mutation rate. Another hypothesis is that RdDM mutants have increased TE activity (Baduel et al. 2021; Zhang et al. 2023), and this leads to genomic instability or dedication of repair factors to TE damage. However, TE activity is reportedly higher in *ddm1* than in *nrpd1* (Baduel et al. 2021; Zhang et al. 2023), so we would expect *ddm1* to have a higher mutation rate than *nrpd1* under this hypothesis. Instead, RdDM may have a function in DNA repair that is not restricted to RdDM methylation targets. This hypothesis may be supported by the observation that double strand break repair by single-strand annealing is less efficient in RdDM mutants (Wei et al. 2012). Future work could assess whether DNA damage rates are higher or DNA repair activity is lower in RdDM mutants.

### Conclusions

Through our analysis of somatic mutations in DNA repair and chromatin-related mutants, we draw conclusions on the functions of DNA repair genes, the regional bias of DNA repair pathways, and the mutational consequences of perturbing chromatin state. We find that the mismatch repair activity of animal MutSα appears to be partitioned between MutSα and MutSγ in *Arabidopsis*. MMR and NER are less efficient in TE regions, and MMR is more efficient in accessible chromatin regions. Lastly, perturbation of most histone modification/variants had no measurable effect on mutation rate. Instead, mutation rate was increased in *h2a.w.7*, *ddm1*, and RdDM mutants.

## Methods

### Plant material

Genotypes *arp* (SALK_021478) (Córdoba-Cañero et al. 2011), *polλ-1* (SALK_075391C) (Amoroso et al. 2011), *msh2-1* (SALK_002708) (Hoffman et al. 2004), *mshc* (SALK_037557) (Gonzalez and Spampinato 2020), *uvh3* /*gl1*(CS3820) (Liu et al. 2001), *chr8-1* (SALK_000799) (Al Khateeb et al. 2019), *rad7a-1* (SALK_095626C) (Lahari et al. 2018), *rad5a-2* (SALK_047150C) (Chen et al. 2008), *ku80* (SALK_016627C) (Jia et al. 2012), *parp2-3* (SALK_140400C) (Gu et al. 2019), *sdg8-2* (SALK_026442) (Zhao et al. 2005), *h2a.w.7* (GABI_149G05) (Lorković et al. 2017), and *ddm1* (SALK_000590) (Qüesta et al. 2013) were obtained from the Arabidopsis Biological Resource Center; *clf* (SALK_139371) was from Franziska Turck (Farrona et al. 2011); *h2a.x.3/5* (SALK_012255/SAIL_382_B11) (Lorković et al. 2017) from Frédéric Berger; ros1-7 (Williams et al. 2015) from Mary Gehring; *suvh4/5/6* (SALK_044606/*suvh5-2*/*suvh6-1*) (Inagaki et al. 2010), *FRI pie1-1* (Noh and Amasino 2003), *met1-3* epiRIL-12 (Reinders et al. 2009), *nrpe1-11* (SALK_029919) (Pontes et al. 2006), and *nrpd1*-4 (SALK_083051) (Herr et al. 2005) were from existing lab stocks. All genotypes were in the Col-0 background except for *FRI pie1*, which was in Ws.

All plants were grown in Sungro soil containing Osmocote fertilizer at 21°C under 16 hours of light from a mix of Philips F96T8/TL841 PLUS and Sylvania Octron Eco fluorescent lamps.

Plants were harvested at the opening of the first flower; flowers and flower buds were removed and the remaining shoot was flash frozen.

### NanoSeq

Each frozen tissue sample was ground with a mortar and pestle. DNA was isolated with a DNeasy Plant Mini Kit (Qiagen, 69106) following manufacturer’s instructions. DNA was fragmented to 150bp using a Covaris E220 Evolution Focused-ultrasonicator by the Georgia Genomic and Bioinformatics Core. 200ng of fragmented DNA was size selected using in-house AMPure XP magnetic beads (0.79 beads:sample ratio for lower and 2.15 for upper selection). End blunting was performed by incubating in a reaction of 1x S1 nuclease buffer and 10units S1 nuclease (Thermo, EN0321) for 30 min at RT. The reaction was stopped by addition of 3µL EDTA, cleaned up with magnetic beads (1.8 ratio), and eluted in 32µL 10mM TrisHCl (pH 8). A-tailing and nick blocking was performed by first adding 5µL 10x *E. coli* DNA ligase buffer (NEB, M0205S), 4.5µL 10mM ATP, and 1uL 10units/µL T4 Polynucleotide Kinase (NEB, M0201S) and incubating at 37°C for 30 min. Samples were then placed on ice, 1uL 10unit/µL *E. coli* DNA ligase (NEB, M0205S) was added, and samples were incubated at 16°C for 30 min. A mix of 1.75µL NF H_2_O, 0.25µL 10mM dATP, 0.5µL ddCTP, 0.5µL ddGTP, 0.5µL ddTTP (Cytiva, 27204501), and 3µL 5units/µL Klenow fragment (3’ -> 5’ exo-) (NEB, M0212S) was added and incubated at 37°C for 30 min. Samples were cleaned up with magnetic beads (1.8 ratio) and 2µL 8µM iTruSeq adapter stubs (sequence below) were added. Adapter ligation was performed in a 25µL reaction of 1x T4 DNA ligase buffer and 20units/µL T4 DNA ligase (NEB, M0202S). Samples were incubated at 16°C for 16 hours and cleaned up with two rounds of magnetic beads (1.4 ratio).

DNA concentration was measured by Qubit, and a portion of each library was diluted to 0.1ng/µL. Three dilutions were used to make a 0.0125ng/µL dilution. These dilutions and three 0.1ng/µL dilutions of a previously sequenced library were run on qPCR in triplicate 10µL reactions of 1x Luna qPCR universal master mix (NEB, M3003S), 1.5µM Illumina i5 indexed primer, and 1.5µM Illumina i7 indexed primer (sequences below). The qPCR program was 95°C (60s), 35x (95°C (15s), 60°C (30s)). Cq values of new and previously sequenced libraries were compared to determine the optimal volume of each sample to use per million read pairs sequenced, such that the amount of callable coverage is maximized. An optimal volume of sample for the amount of sequencing planned was used for further steps. Samples were split into multiple tubes to reduce the chance of DNA molecules with the same alignment start and end position being present in a single library.

Samples were amplified in a 50µL reaction of 0.33µM Illumina i5 indexed primer, 0.33µM Illumina i7 indexed primer, 200µM dNTPs, 1x Phusion HF Buffer, and 0.04units/µL Phusion HF DNA Polymerase (NEB, M0530S). PCR program was 98°C (60s), 2x (98°C (15s), 50°C (120s), 72°C (15s)), 11x (98°C (15s), 60°C (30s), 72°C (15s)). Samples were cleaned up with magnetic beads (1.4 ratio) and sequenced on the NovaSeq X series platform 25B flow cell to generate 150bp paired end reads. Genotypes were sequenced to a median depth of 1.72B read pairs (**Supplemental Table S3**). iTruSeq stub sequences: ACACTCTTTCCCTACACGACGCTCTTCCGATCT and /5phos/GATCGGAAGAGCACACGTCTGAACTCCAGTCAC Illumina i5 and i7 indexed primers: AATGATACGGCGACCACCGAGATCTACACNNNNNNNNACACTCTTTCCCTAC and CAAGCAGAAGACGGCATACGAGATNNNNNNNNGTGACTGGAGTTCAG where Ns are the sample index

### Filtering and alignment of reads

Sequencing reads were trimmed and filtered with fastp using default parameters (Chen 2023). Reads were aligned to the TAIR10 reference genome using Bowtie2 with -X 800 (Lamesch et al. 2011; Langmead and Salzberg 2012). Optical duplicates were then marked using SAMtools fixmate -m, Sambamba sort, and SAMtools markdup -d 2500 (Tarasov et al. 2015; Danecek et al. 2021). Optical duplicates, non-concordantly mapped, and ambiguously mapped reads were filtered using sambamba view --filter mapping_quality ≥ 1 and proper_pair and ([dt] == null or [dt] != ‘SQ’). Replicate libraries generated from the same plant were then merged using SAMtools merge -r to produce the final filtered BAM files.

67% of read pairs were retained after trimming, alignment, optical duplicate removal, and filtering. The median number of callable molecules per retained read pair was 0.073, and the median callable bps per retained read pair was 6.36 (**Supplemental Table S3**, **Supplemental Fig. S13**).

### Filtering somatic mutations

Somatic mutations were identified as described in Meyer et al. (2025). A set of unfiltered variants were identified within each NanoSeq molecule (a set of PCR duplicates or “read bundle”) using a custom script. These variants were then filtered by passing the following requirements: ≥24% of reads in the top strand duplicates and ≥24% of reads in the bottom strand duplicates support the variant; ≥6 duplicate reads cover the variant position; ≥2 top strand duplicates and ≥2 bottom strand duplicates cover the variant; ≥4 duplicate reads support the variant with a BQ >30 at the variant base (if SNV); ≥10 average MQ of supporting reads; ≤0.21 mismatches/bp between the variant and each fragment end (this removes all variants <5bp from a fragment end); variant is ≥6bp from fragment ends (applied to indels only); variant position is ≥0.7 percentile in total read coverage across the eight original WT libraries (i.e. remove genomic positions with low read coverage); variant position is not the start/end of a poly-A nor poly-T repeat of length ≥8; variant position is not the start/end of a dinucleotide nor trinucleotide repeat of ≥5 repeating units; variant is supported by the majority of reads in ≤3 molecules across the eight original WT libraries; variant is in a fragment of length ≤300bp; there are ≤4bp between any 2 variants in the same molecule passing all previous filters (this filter was not applied to *FRI pie1*, as it has a large number fixed variants); ≥76% of reads in the top strand duplicates and ≥76% of reads in the bottom strand duplicates support the variant. Diagrams of each filter and the methods used to determine optimal thresholds for each filter were previously reported in Meyer et al. (2025).

The frequency of each mutation in the sample was calculated as the fraction of molecules covering the variant site which had >76% of reads supporting the variant. Only molecules with at least two reads covering the variant site were considered. Variants identified in more than one plant or at a frequency >0.35 were discarded.

For the *polλ* samples, we detected trace levels of contamination from Ler-0 DNA, so any variants found in >4 molecules of a previously generated Ler-0 NanoSeq library (SRR33007844) were removed. For the *FRI pie1* samples, the Ws background introduced false positives which were removed by discarding variants present in >0 molecules of a sibling library, within 10bp of an indel, or at a site with <20 callable coverage across *FRI pie1* samples.

A list of all somatic mutations identified can be found in **Supplemental Table S4** and at https://zenodo.org/records/21249613?preview=1Ctoken=eyJhbGciOiJIUzUxMiJ9.eyJpZCI6ImQ4OTk4YTExLWE3OTAtNDg5NS04NTM4LWU1ZjBlZTdkMjhiZCIsImRhdGEiOnt9LCJyYW5kb20iOiIzNGNkYjI3ZmNhNGQxMDA4ZTE2NjZjODhiNTQwZDQzOSJ9.U_h0YlfAtRiW2FctqhNkmA76cKk0WRiHgIv1l3vRiTFtOj8U38pq8odsz23fLf4LhE2LnXcNmCGsVfJ5gIPq_A.

### Calculation of callable coverage and somatic mutation rate

Callable coverage was calculated per site as the number of opportunities to observe a somatic mutation that could pass all filters. Each molecule was considered “callable” if it had ≥1 bottom strand, ≥1 top strand, and ≥3 total read pairs, as well as average MQ ≥10 and fragment length ≤300bp. Then, each base within a callable molecule contributed to callable coverage if it overlapped ≥2 reads from the bottom strand, ≥2 reads from the top strand, and ≥6 total reads, was ≥5bp from a molecule end, and was not in any blacklisted regions of the genome. Blacklisted regions were those with <0.7 percentile total read coverage across the eight original WT libraries, ≥8bp poly-A and poly-T repeats, and ≥5 length di/trinucleotide repeats. For the *FRI pie1* samples, sites within 10bp of an indel or <20 callable coverage across *FRI pie1* samples were also blacklisted to match the filters unique to those samples.

To calculate the overall somatic mutation rate of a sample, the callable coverage of all sites in the nuclear genome was summed and divided by a predicted rate of molecule conflicts. The conflict rate represents the chance any given molecule in the library has another molecule with the same alignment start and end site, which may preclude mutation identification in that molecule. Conflict rate was estimated by randomly selecting half of the molecules from two libraries of the same plant and calculating a “half” conflict rate as the fraction of selected molecules which conflict with another selected molecule. The full conflict rate was then calculated as 1 – (1 – half conflict rate)^2^. For samples with only one library per plant, the conflict rate was imputed using the other samples assuming a linear relationship between conflict rate and molecules in the library (**Supplemental Fig. S14**). Thus, the overall somatic mutation rate was calculated as the number of mutations / (callable coverage * (1 – conflict rate)). The same formula was used to calculate the somatic mutation rate of a genomic region by only considering mutations and callable covering within the region.

The callable coverage per sample can be found in **Supplemental Table S3**, and wiggle track files of callable coverage per genomic site can be found at https://zenodo.org/records/21249613?preview=1Ctoken=eyJhbGciOiJIUzUxMiJ9.eyJpZCI6ImQ4OTk4YTExLWE3OTAtNDg5NS04NTM4LWU1ZjBlZTdkMjhiZCIsImRhdGEiOnt9LCJyYW5kb20iOiIzNGNkYjI3ZmNhNGQxMDA4ZTE2NjZjODhiNTQwZDQzOSJ9.U_h0YlfAtRiW2FctqhNkmA76cKk0WRiHgIv1l3vRiTFtOj8U38pq8odsz23fLf4LhE2LnXcNmCGsVfJ5gIPq_A.

### Calculation of mutation spectra

Mutation rate was calculated for each SNV type, 3bp context, insertions, and deletions as the count of that mutation type divided by the callable coverage of sites which could harbor that mutation (*e.g.* coverage of all C·G pairs for C>T mutations, coverage of all ACA sites for ACA>AAA mutations, and coverage of all sites for indels). This mutation rate was then divided by the genome-wide average mutation rate for that sample to yield the relative mutation rate in **Supplemental Figure S3B**. Raw mutation counts per SNV and raw mutation rates per SNV can be found in **Supplemental Table S5**.

### Gene, TE, and “other” regions

Gene, TE, and “other” regions of the genome were defined by first labeling all sites within a TE of the Panda and Slotkin (2020) annotation as “TE”. All sites within mRNA elements of the TAIR10 annotations were labeled as “gene” unless already labeled as “TE”. All remaining sites in the genome were labeled as “other”.

### Accessible chromatin regions

Two replicates of Col-0 ATAC-seq and a gDNA control were downloaded from NCBI SRA PRJNA527732 (Lu et al. 2019). Sequencing reads were trimmed and filtered with fastp using default parameters (Chen 2023). Reads were aligned to the TAIR10 reference genome using Bowtie2 with -X 800 (Langmead and Salzberg 2012). Mapped reads were converted to a BED file of Tn5 insertions using BEDTools, where the positions 5bp after the start of each forward mapped read and 6bp before the end of each reverse mapped read were labeled as insertion sites. Peaks were called for each replicate using MACS2 callpeak -g 1.1e8 -q 0.7 --nomodel --extsize 200 --shift −100 and the gDNA as a control (Zhang et al. 2008). The union of peaks in the two replicates was used as the final accessible chromatin region set. ACR counts and genome browser screenshots can be found in **Supplemental Table S6** and **Supplemental Figure S8**.

### DNA methylation

Methylated cytosines in WT and *met1* epiRIL 12 were identified using whole genome bisulfite sequencing data from SRR2922654 and SRR2922657 (Bewick et al. 2016). Reads were trimmed and filtered with fastp using default parameters (Chen 2023). Methylation files were generated using methylpy --trim-reads False --min-mapq 10 --merge-by-max-mapq --unmethylated-control “ChrC” --binom-test True (Schultz et al. 2015). Cytosines with a “1” in the methylated column were labeled as methylated and those with a “0” as unmethylated.

To find sites which lost methylation in the *met1* epiRIL, we first manually identified blocks of the genome with a high rate of exonic CG sites methylated in the WT but not the epiRIL (**Supplemental Fig. S15**). Only the CG sites within these blocks, methylated in WT, and unmethylated in the *met1* epiRIL were labeled as methylation loss sites.

To identify differentially methylated regions in *ddm1*, *nrpe1*, and *nrpd1*, we processed bisulfite sequencing data from PRJNA172021 (Stroud et al. 2013) as described above.

Differentially methylated regions were called using methylpy DMRfind --min-num-dms 10 -- sig-cutoff 0.3 –dmr-max-dist 500 on the two WT controls and the *ddm1*, *nrpe1*, or *nrpd1* sample (Schultz et al. 2015). For *nrpe1* and *nrpd1*, --mc-type was set to CHN (i.e. only non-CG sites are considered), whereas for *ddm1* it was set to CNN (all contexts are considered). Only differentially methylated regions with a >0.1 absolute difference in methylation rate between the mutant and the average of the WT samples were retained (e.g. a region with a 20% methylation rate in WT and <10% in *ddm1* will be labeled hypomethylated). DMR counts and genome browser screenshots can be found in **Supplemental Table S6** and **Supplemental Figure S10**.

### ChIP-seq

WT ChIP-seq data for H3K36me3, H3K27me3, and H2A.Z was sourced from PRJNA527732 (Lu et al. 2019); H3K9me2 from PRJNA302602 (Bewick et al. 2016); H2A.X from PRJNA689609 (Osakabe et al. 2021); H2A.W.7 from PRJNA377526 (Lorković et al. 2017); and NRPE1 from PRJNA390422 (Liu et al. 2018). WT and *sdg8* H3K36me3 ChIP-seq data for identifying regions which lose H3K36me3 in *sdg8* from PRJNA1068029 (Yao et al. 2025).

Reads were trimmed and filtered with default fastp parameters and aligned to the reference genome with Bowtie2 -X 800 (Langmead and Salzberg 2012; Chen 2023). Peaks were called using MACS2 with different parameters for each sample (**Supplemental Table S6**). Peak call counts and genome browser screenshots can be found in **Supplemental Table S6** and **Supplemental Figure S8**.

### RNA-seq

RNA-seq data of three WT and three *nrpe1* plants was obtained from PRJNA915364 (Li et al. 2023). Reads were trimmed and filtered with default fastp parameters and aligned to the reference genome with STAR --quantMode GeneCounts (Dobin et al. 2013; Chen 2023).

Differentially expressed genes were determined with DESeq2 (**Supplemental Table S2**) (Love et al. 2014). Gene ontology enrichment was performed on the upregulated and downregulated gene sets using PANTHER on the GO biological process dataset (Mi et al. 2019; Thomas et al. 2022).

### Statistical tests

All statistical tests reported in the text and in figures were two-tailed Student’s t-tests performed with the SciPy package (Virtanen et al. 2020). The Holm method was used to correct for multiple hypothesis testing by considering the number of tests performed within the figure panel.

### Software availability

The data processing pipelines, custom python scripts, and python code for figure making are available at https://github.com/cullanm/repair-and-chromatin-mutant-nanoseq.

## Data access

All NanoSeq sequencing data generated in this study have been submitted to the NCBI BioProject database (https://www.ncbi.nlm.nih.gov/bioproject/) under accession number PRJNA1468256 (https://dataview.ncbi.nlm.nih.gov/object/PRJNA1468256?reviewer=u112n6l0t7v10ttsvmu f6aqubk). A list of somatic mutations can be found in **Supplemental Table S4**. Somatic mutations, callable coverage per genomic position, DNA methylation calls, DMRs, ChIP-seq peaks, and RNA-seq counts can be found at https://zenodo.org/records/21249613?preview=1Ctoken=eyJhbGciOiJIUzUxMiJ9.eyJpZCI6ImQ4OTk4YTExLWE3OTAtNDg5NS04NTM4LWU1ZjBlZTdkMjhiZCIsImRhdGEiOnt9LCJyYW5kb20iOiIzNGNkYjI3ZmNhNGQxMDA4ZTE2NjZjODhiNTQwZDQzOSJ9.U_h0YlfAtRiW2FctqhNkmA76cKk0WRiHgIv1l3vRiTFtOj8U38pq8odsz23fLf4LhE2LnXcNmCGsVfJ5gIPq_A.

## Competing interest statement

The authors declare no competing interests.

## Acknowledgements

We would like to thank Alexandra Tadros and Zack Lewis for providing advice on the manuscript, as well as Mark Minow and Xiang Li for their feedback throughout the project. Thanks to Franziska Turck, Frédéric Berger, Mary Gehring, and the Arabidopsis Biological Resource Center, whose mutant seed stocks were included in the study. The Georgia Advanced Computing Resource Center provided the computational resources required for data analysis. This research was supported by the National Science Foundation (MCB-2242696) and the University of Georgia Office of Research to R.J.S. as well as the National Institute for General Medical Sciences of the National Institutes of Health to C.A.M. (1T32GM142623).

## Author contributions

C.A.M. and R.J.S. designed the research and wrote the paper. C.A.M. conducted the research and analyzed the data.

## Notes

https://dataview.ncbi.nlm.nih.gov/object/PRJNA1468256?reviewer=u112n6l0t7v10ttsvmuf6aqubk

https://zenodo.org/records/21249613?preview=1&token=eyJhbGciOiJIUzUxMiJ9.eyJpZCI6ImQ4OTk4YTExLWE3OTAtNDg5NS04NTM4LWU1ZjBlZTdkMjhiZCIsImRhdGEiOnt9LCJyYW5kb20iOiIzNGNkYjI3ZmNhNGQxMDA4ZTE2NjZjODhiNTQwZDQzOSJ9.U_h0YlfAtRiW2FctqhNkmA76cKk0WRiHgIv1l3vRiTFtOj8U38pq8odsz23fLf4LhE2LnXcNmCGsVfJ5gIPq_A

https://github.com/cullanm/repair-and-chromatin-mutant-nanoseq

### Competing Interest Statement

The authors have declared no competing interest.

