## Supplemental Figures for "Mutational consequences of perturbing DNA repair and chromatin state in *Arabidopsis*"

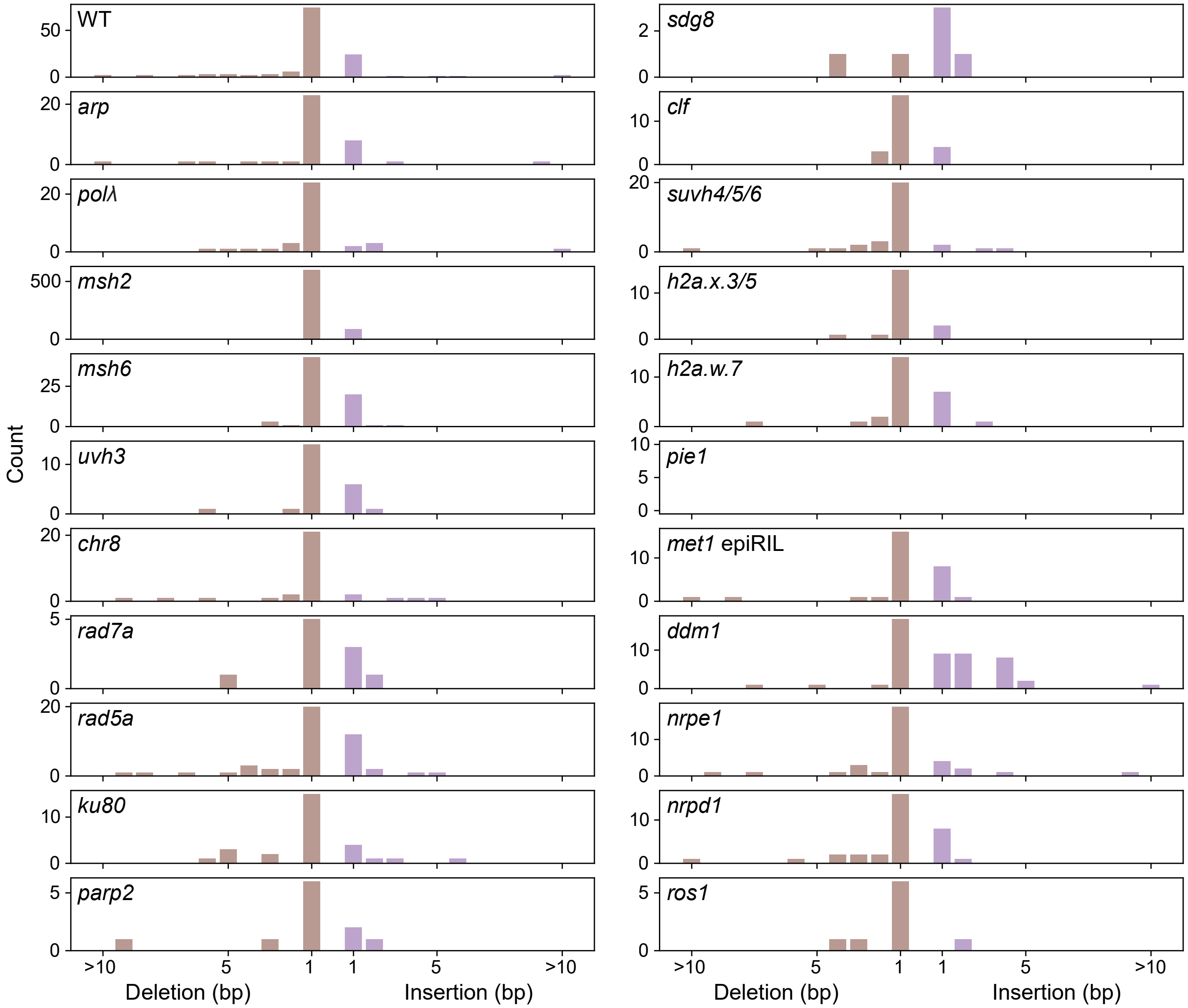

### Supplemental Figure S1. Insertion and deletion size distribution of all genotypes

Histograms of insertion/deletion size. In the WT and most genotypes, deletions are more common than insertions, and this difference in increased in *msh2*. In *ddm1*, insertions are more common, especially those >1bp.

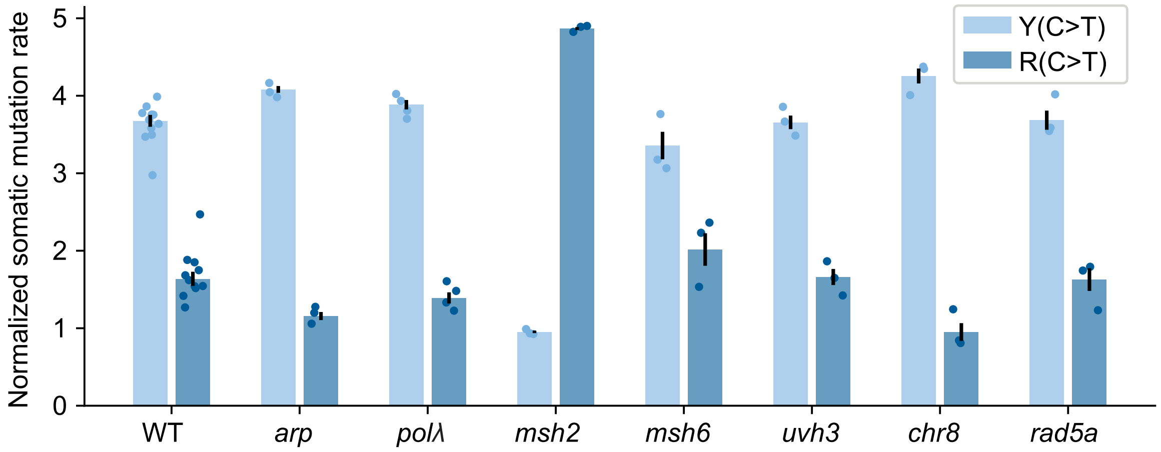

### Supplemental Figure S2. C>T mutation purine/pyrimidine context bias

Normalized somatic mutation rate of C>T mutations at YC and RC sites (Y=C/T, R=A/G). The somatic mutation rate of Y(C>T) and R(C>T) SNVs was divided by the genotype's overall mutation rate to yield a normalized somatic mutation rate for both SNV types. In most genotypes, mutation rate is higher at YC sites, perhaps due to cyclobutane pyrimidine dimer formation. This bias is reversed in *msh2*, suggesting polymerase errors or other MMR repaired damage is more frequent at RC sites.

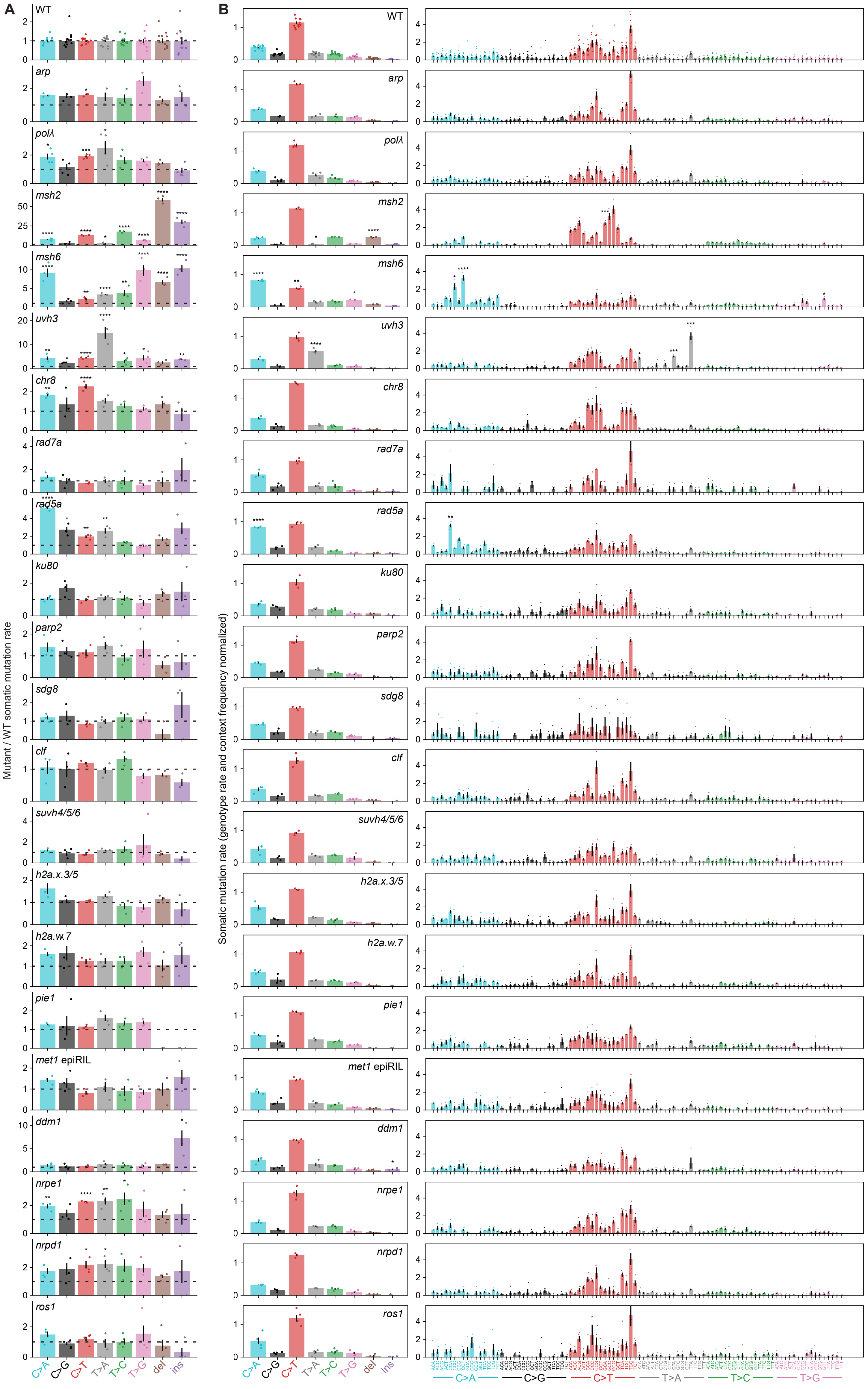

### Supplemental Figure S3. Mutation spectra of all genotypes

**A**, Mutant divided by WT mutation rate of each SNV/indel type. The dotted line indicates a value of one (WT mutation rate). **B**, Normalized somatic mutation rate of each SNV/indel type. The somatic mutation rate of each SNV was divided by the genotype's overall mutation rate to yield a context frequency normalized somatic mutation rate. Mutation spectra were then subdivided by 3bp sequence context (right). See Supplemental Table S5 for raw counts. *msh2* has a greater rate of R(C>T) than Y(C>T) mutations, and *uvh3* has a high rate of T>A at ATA, GTA, and TTA sites. **A&B**, Dots represent individual plants. Significance labels indicate the value differs from WT (Holm-adjusted t-test). *=p<0.05, **=p<0.01, ***=p<0.001, ****=p<0.0001. Error bars=±1 SEM.

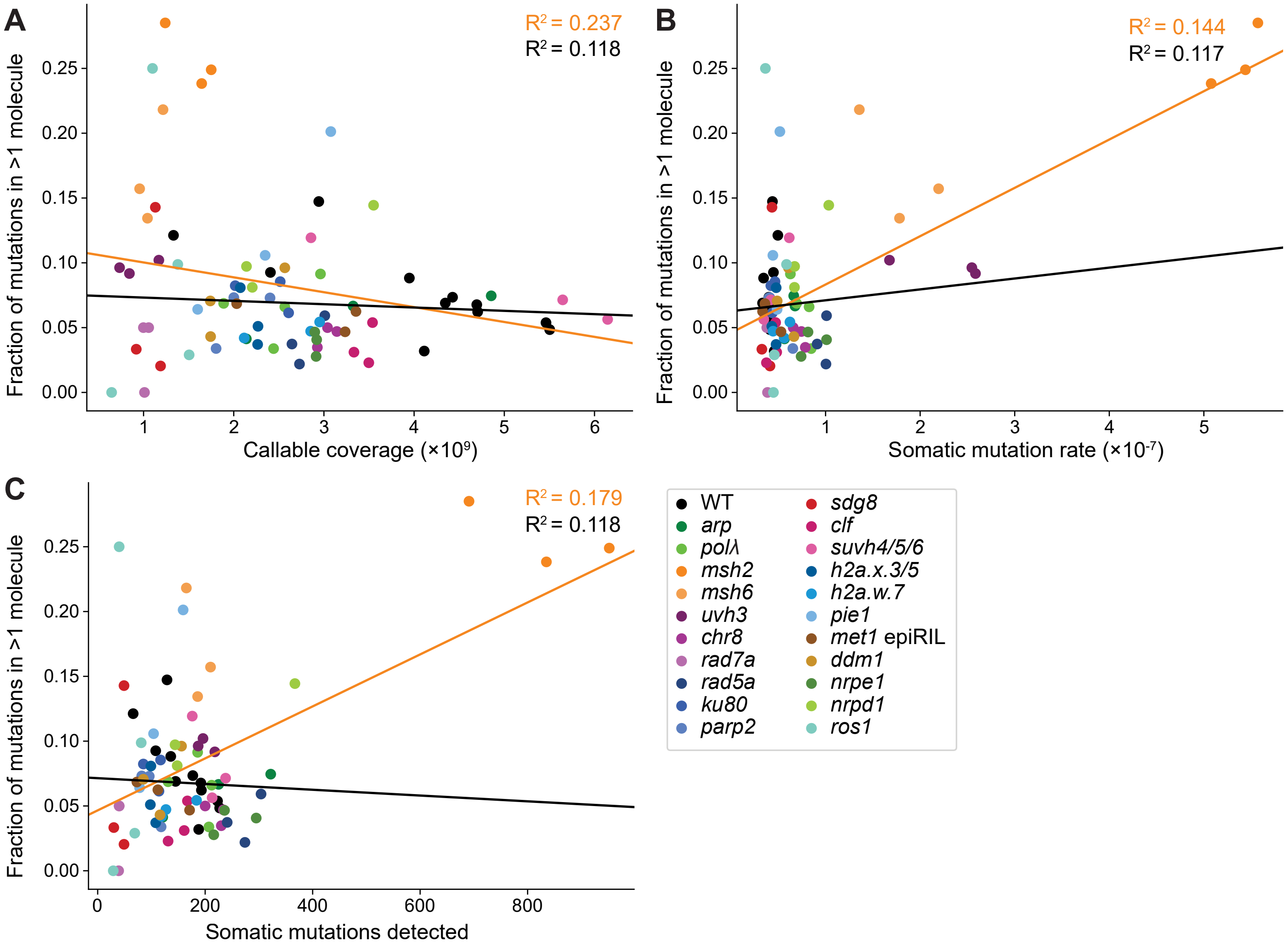

### Supplemental Figure S4. Technical factors do not explain the large proportion of high abundance mutations in *msh2*

Fraction of mutations detected in >1 molecule plotted against the amount of callable coverage (**A**), mutation rate (**B**), and number of mutations detected (**C**) in each NanoSeq library. Orange lines indicate the least squares linear regression of all dots, whereas black lines ignore the *msh2* and *msh6* samples. While mutation rate and number do positively correlate with high abundance mutations, these factors explain only a small portion of the variance ($R^2$), and the correlation disappears when *msh2* and *msh6* are not considered.

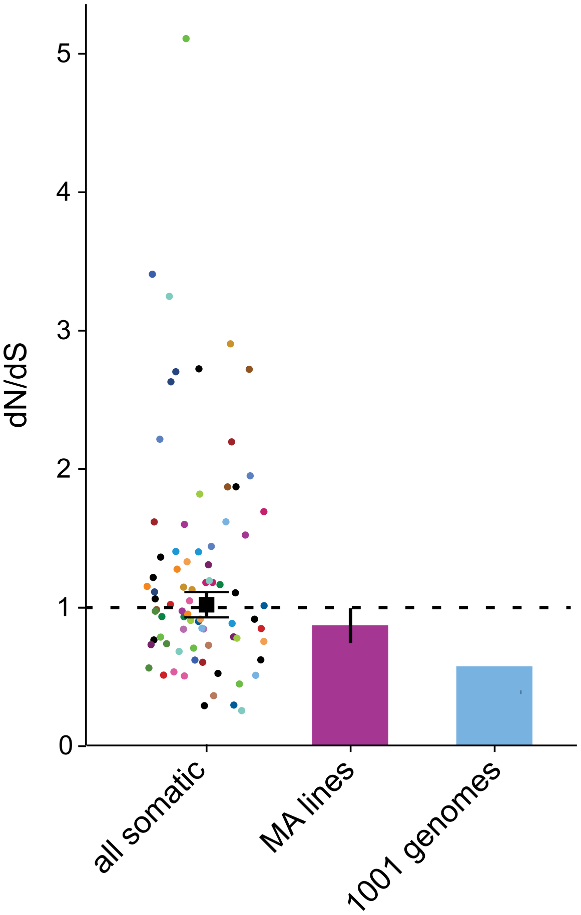

### Supplemental Figure S5. No evidence of selection in somatic mutations

Mutation/polymorphism rate at non-synonymous sites divided by synonymous sites. Dots indicate the somatic mutation rate dN/dS of every NanoSeq sample in this study. The “MA lines” and “1001 genomes” values were calculated in Meyer et al. (2025). The MA line data was originally from (Weng et al. 2019), and non-synonymous/synonymous mutation rates were calculated from each MA line by dividing the number of mutations by the number of sites with at least one callable coverage in the WT NanoSeq libraries. The 1001 genomes data was originally from Alonso-Blanco et al. (2016), and non-synonymous/synonymous polymorphism rate for the entire dataset was calculated as the number of polymorphisms divided by the number of sites with at least one callable coverage in the WT NanoSeq libraries. Error bars=±1 SEM.

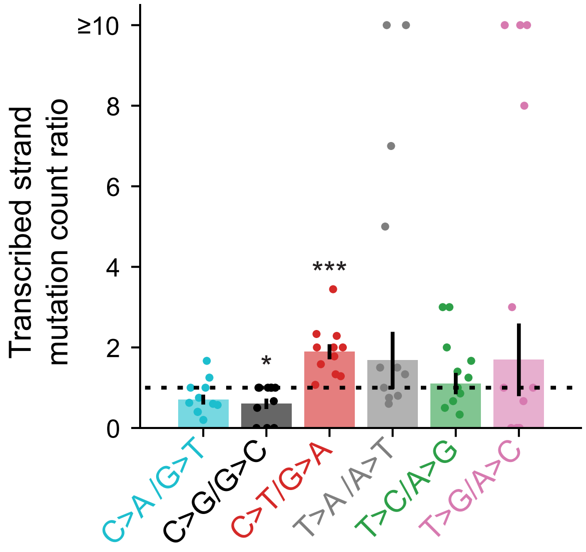

### Supplemental Figure S6. SNV strand asymmetries in WT

Number of mutations on where the C/T is on the non-transcribed strand divided by the transcribed strand for each SNV type. Significance labels indicate whether the average ratio differs from a value of one (Holm-adjusted t-test). *=p<0.05, **=p<0.01, ***=p<0.001, ****=p<0.0001. Error bars=±1 SEM. There are significantly more C>G mutations when the C is on the transcribed strand and significantly more C>T mutations when the C is on the non-transcribed strand.

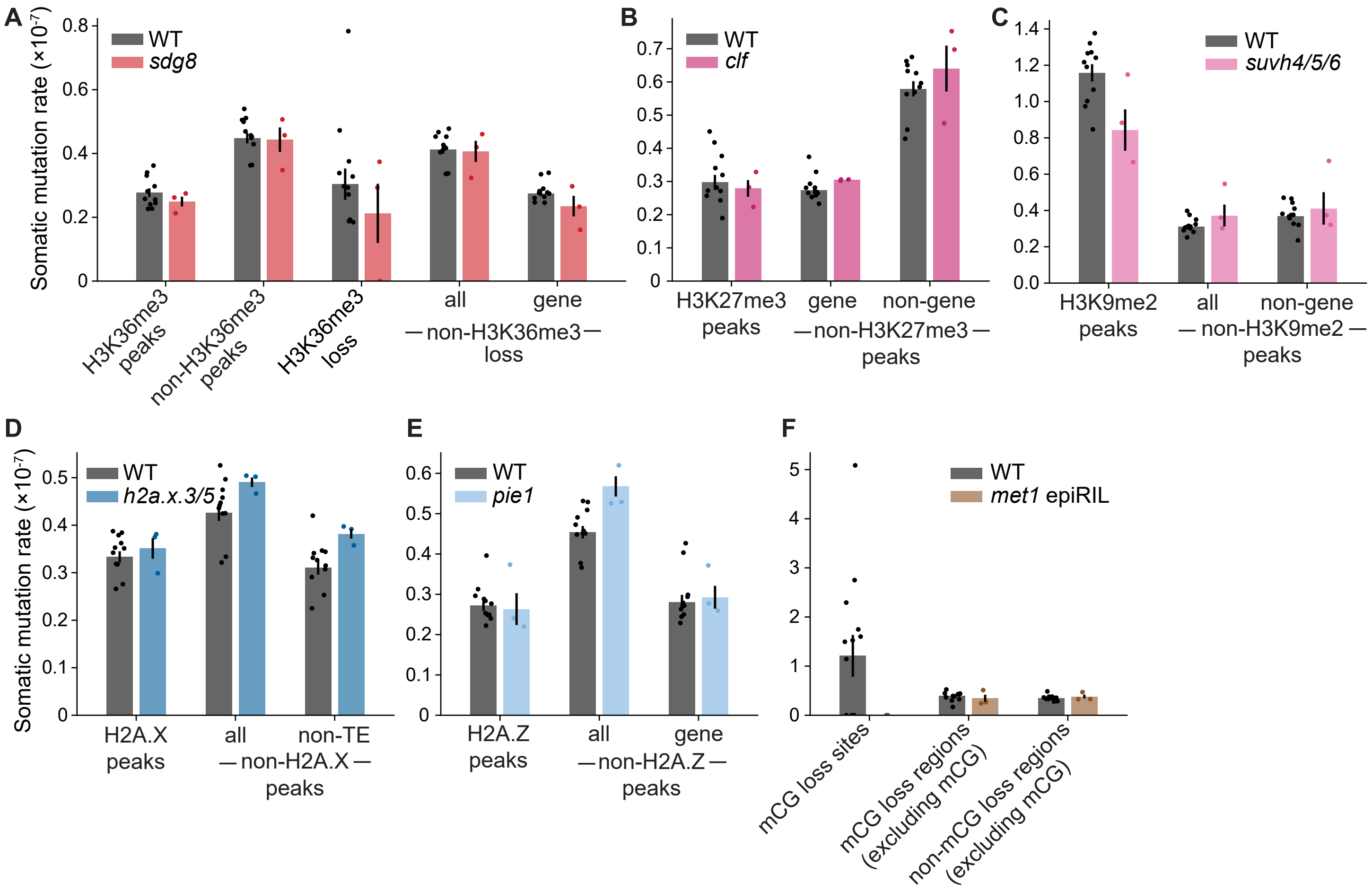

### Supplemental Figure S7. Mutation rates of chromatin mutants in ChIP-seq peaks

**A**, Somatic mutation rate of WT and *sdg8* in H3K36me3 ChIP-seq peaks and regions depleted for H3K36me3 ChIP-seq signal in *sdg8*. **B**, Somatic mutation rate of WT and *clf* in H3K27me3 ChIP-seq peaks. **C**, Somatic mutation rate of WT and *suvh4/5/6* in H3K9me2 ChIP-seq peaks. **D**, Somatic mutation rate of WT and *h2a.x.3/5* in H2A.X ChIP-seq peaks. **E**, Somatic mutation rate of WT and *h2a.w.7* in H2A.W.7 ChIP-seq peaks. **F**, Somatic mutation rate of WT and *pie1* in H2A.Z ChIP-seq peaks. **G**, Somatic mutation rate of WT and the *met1* epiRIL at CG sites which lose methylation in the *met1* epiRIL, genomic windows which lost mCG in the *met1* epiRIL (excluding the mCG sites themselves), and genomic windows which retained mCG in the *met1* epiRIL (excluding the mCG sites). **A-G**, Dots represent individual plants. No comparisons between WT and the mutant within the same region were statistically significant (Holm-adjusted t-tests). Error bars=±1 SEM. We observe no statistically significant differences in mutation rate within ChIP-seq peaks when the histone modification/variant is perturbed.

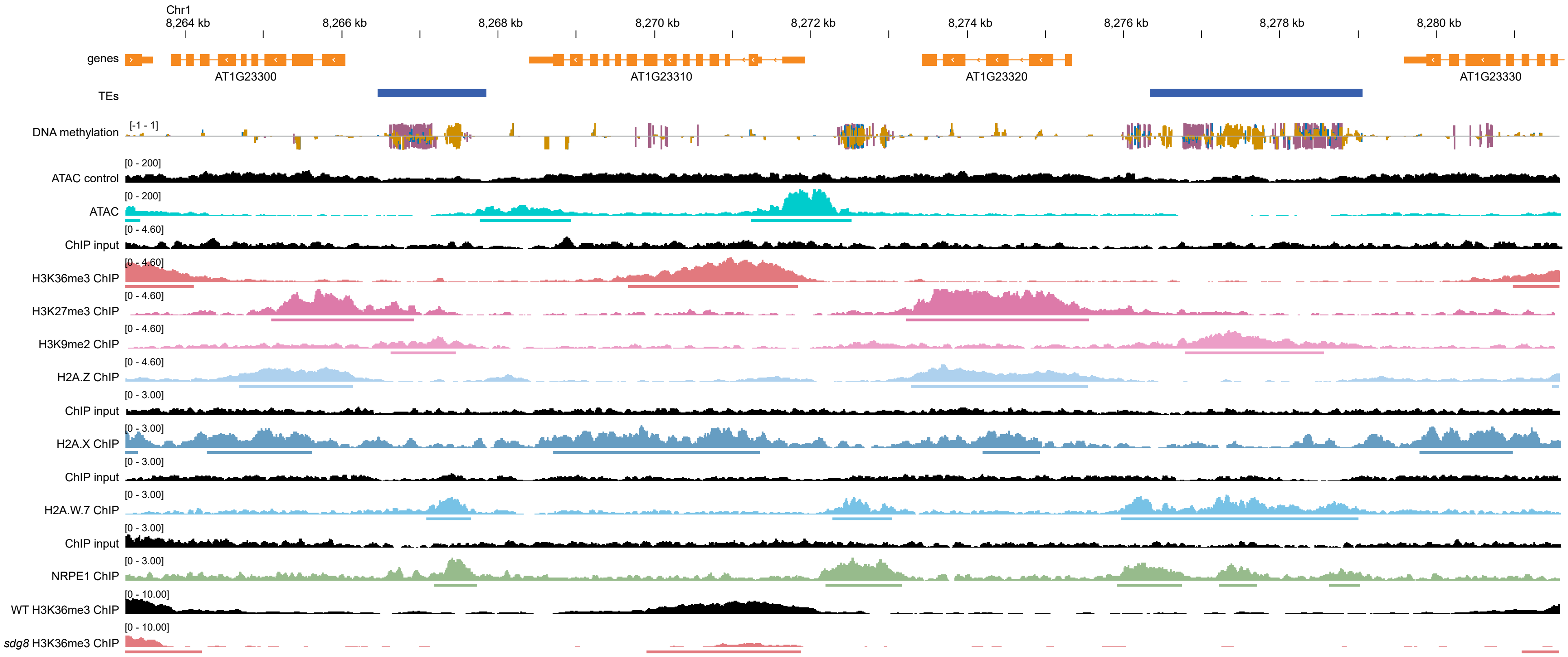

### Supplemental Figure S8. Genome Browser view of ATAC-seq and ChIP-seq data

Genome browser tracks of TAIR10 annotated genes, Panda and Slotkin (2020) annotated TEs, WT DNA methylation, ATAC-seq, histone modification ChIP-seq, histone variant ChIP-seq, NRPE1 ChIP-seq, and WT and *sdg8* H3K36me3 ChIP-seq. Controls are colored black, representing ATAC-seq on gDNA, ChIP-seq input (H3K36me3, H3K27me3, H3K9me2, and H2A.Z all used the same input), and WT H3K36me3 ChIP-seq (for the *sdg8* H3K36me3 ChIP-seq). Peak calls are shown as lines below the enrichment tracks. For the *sdg8* H3K36me3 ChIP-seq, peak calls represent regions with diminished H3K36me3 signal compared to WT.

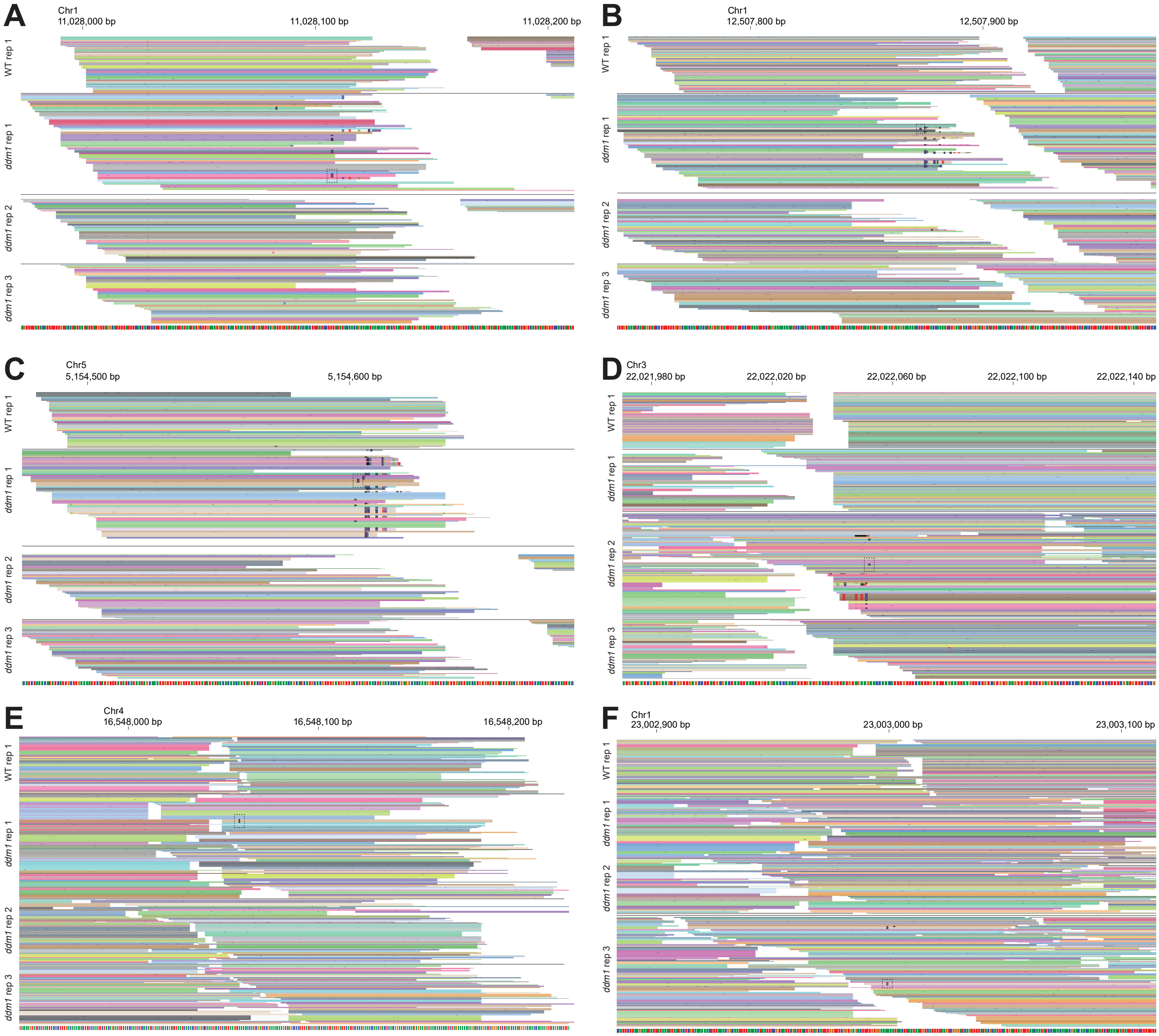

### Supplemental Figure S9. Potential structural variants in *ddm1*

**A-G**, Genome browser views of insertions identified in *ddm1* which may suggest structural variants (**A-D**) and those which do not suggest structural variants (**E&F**). Reads from one WT sample and all 3 *ddm1* samples are shown in each panel. Each line represents one sequencing read. Reads of the same color are PCR duplicates of a single NanoSeq molecule. Bases which differ from the reference are colored green (A), blue (C), yellow (G), or red (T) while insertions are colored purple with a black outline. The insertion called as a somatic mutation is enclosed in a box of dotted lines.

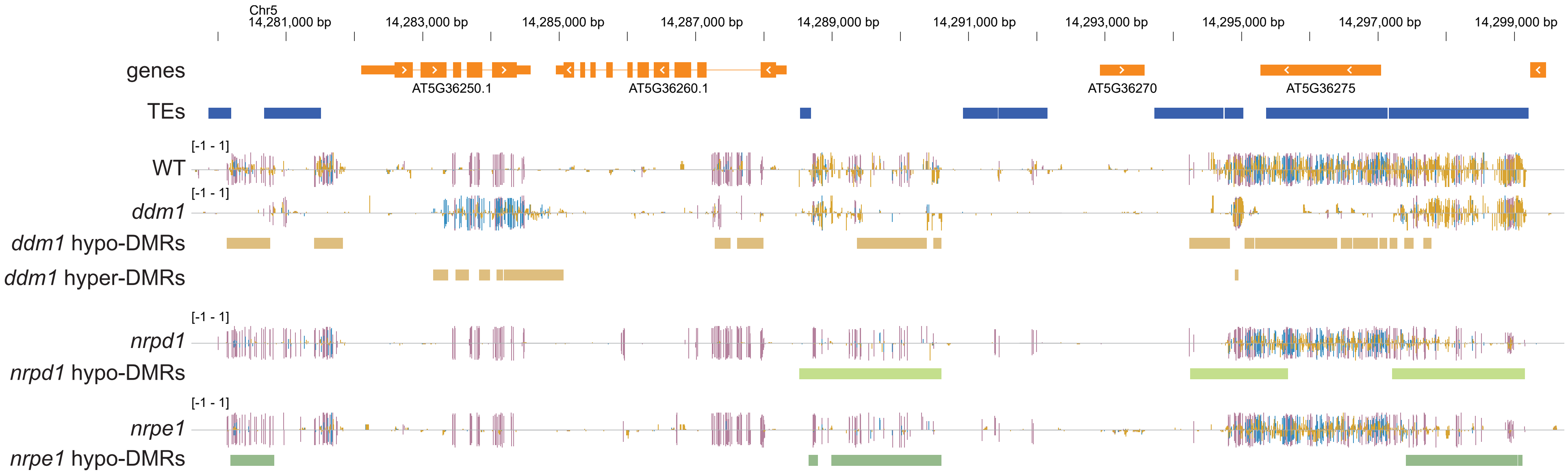

### Supplemental Figure S10. Genome browser view of DMRs

Genome browser view of WT, *ddm1*, *nrpd1*, and *nrpe1* DNA methylation and differentially methylated regions (DMRs). Negative DNA methylation values represent cytosines on the reverse strand. Purple=mCG, blue=mCHG, yellow=mCHH (H=A/C/T). Genes are from the TAIR10 annotation while TEs are from Panda and Slotkin (2020)

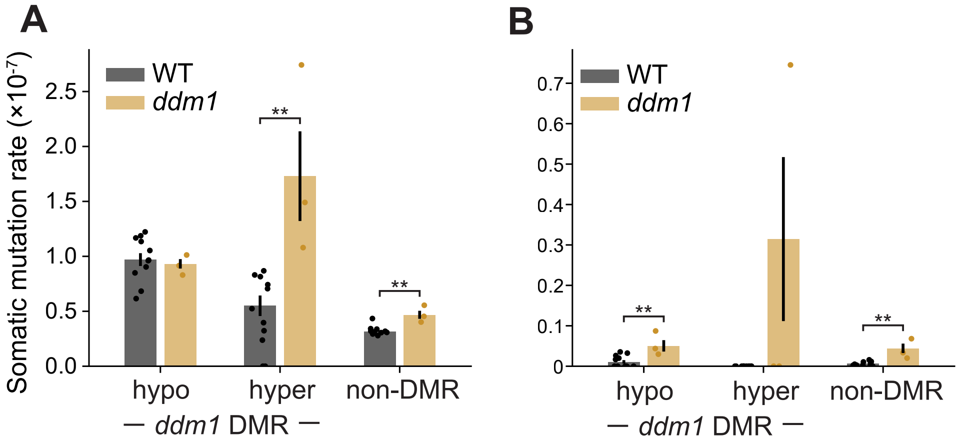

### Supplemental Figure S11. *ddm1* mutation rate when discarding insertions or considering only insertions

**A&B**, Somatic mutation rate of WT and *ddm1* in *ddm1* hypomethylated DMRs, hypermethylated DMRs, and non-DMRs when removing insertions from the analysis (**A**) or considering only insertions (**B**). Dots represent individual plants. Significance values indicate Holm-adjusted t-tests. *=p<0.05, **=p<0.01, ***=p<0.001, ****=p<0.0001. Error bars=±1 SEM. Mutation rate is elevated in hypermethylated DMRs and non-DMRs even when insertions are not considered.

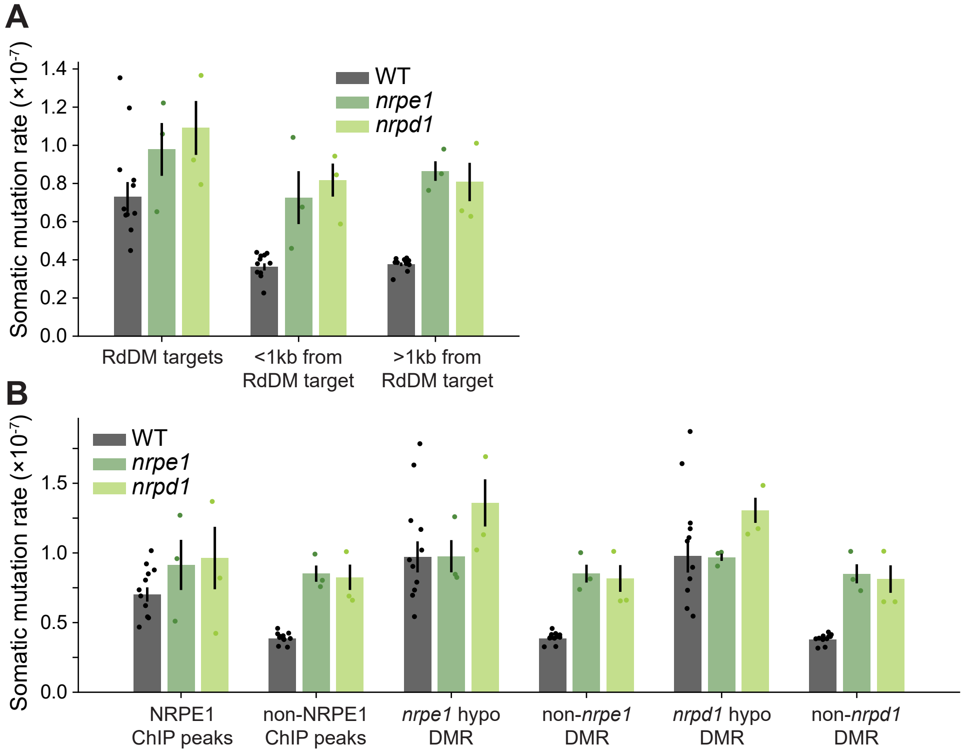

### Supplemental Figure S12. *nrpe1* and *nrpd1* mutation rate near RdDM targets and using other definitions of RdDM target

**A**, Somatic mutation rate of WT, *nrpe1*, and *nrpd1* in RdDM targets, regions within 1kb of an RdDM target, and regions >1kb from an RdDM target. RdDM targets are defined as the union of NRPE1 ChIP-seq peaks, *nrpe1* hypomethylated DMRs, and *nrpd1* hypomethylated DMRs. The mutation rate increase in *nrpe1* and *nrpd1* is not restricted to RdDM targets nor regions near RdDM targets. **B**, Somatic mutation rate of WT, *nrpe1*, and *nrpd1* in NRPE1 ChIP-seq peaks, *nrpe1* DMRs, and *nrpd1* DMRs. Mutation rate is increased both within and outside of RdDM targets in *nrpe1* and *nrpd1*, regardless of which evidence is used to define RdDM targets. **A&B**, Dots represent individual plants. Error bars=±1 SEM.

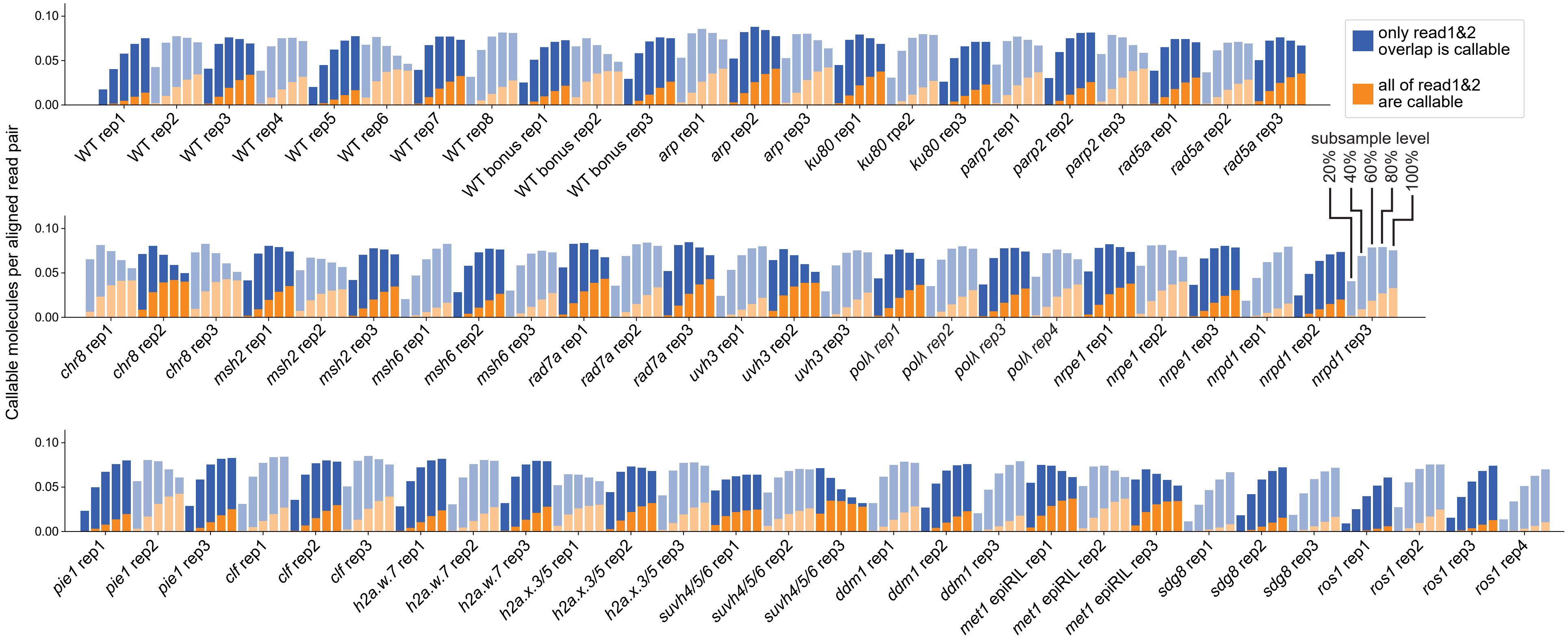

### Supplemental Figure S13. Dilution efficiency of all NanoSeq libraries

Sequencing efficiency of NanoSeq libraries measured as callable molecules per properly aligned read pair. Molecules are considered callable in the read1-read2 overlap if they have ≥1 read pair originating from each strand and ≥3 read pairs in total. Molecules with ≥2 read pairs per strand and ≥6 read pairs total are callable in all of read1 and read2. Libraries were randomly subsampled to various percentages of read pairs to determine whether less sequencing would have increased the sequencing efficiency (i.e. the library was oversequenced). Libraries with peak sequencing efficiency at the 100% subsample level were sequenced optimally.

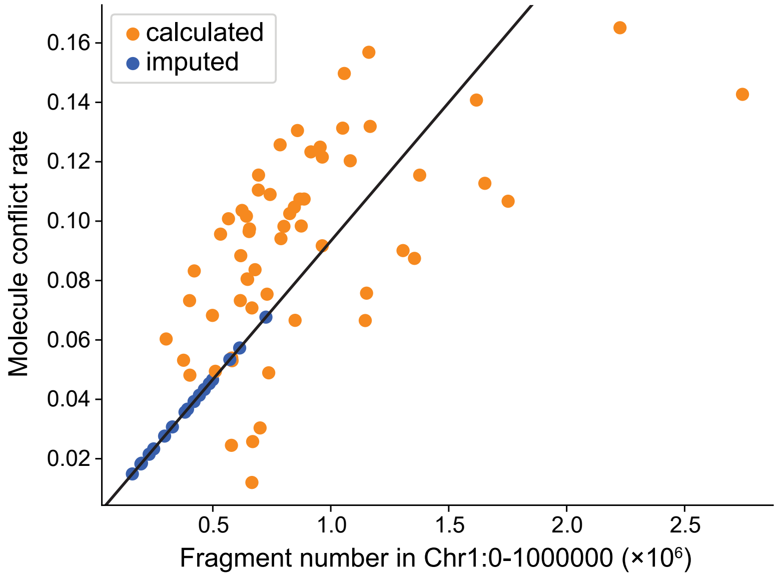

### Supplemental Figure S14. Molecule conflict rate for all NanoSeq libraries

Molecule conflict rate and molecules sequenced in each NanoSeq library. Conflict rate was calculated as the fraction of NanoSeq molecules where another molecule in the library has the same alignment start and end position (Methods). Conflict rate correlates with the number of molecules sequenced, so for libraries which could not have their conflict rate calculated (those with only one split), it was imputed from the line of best fit.

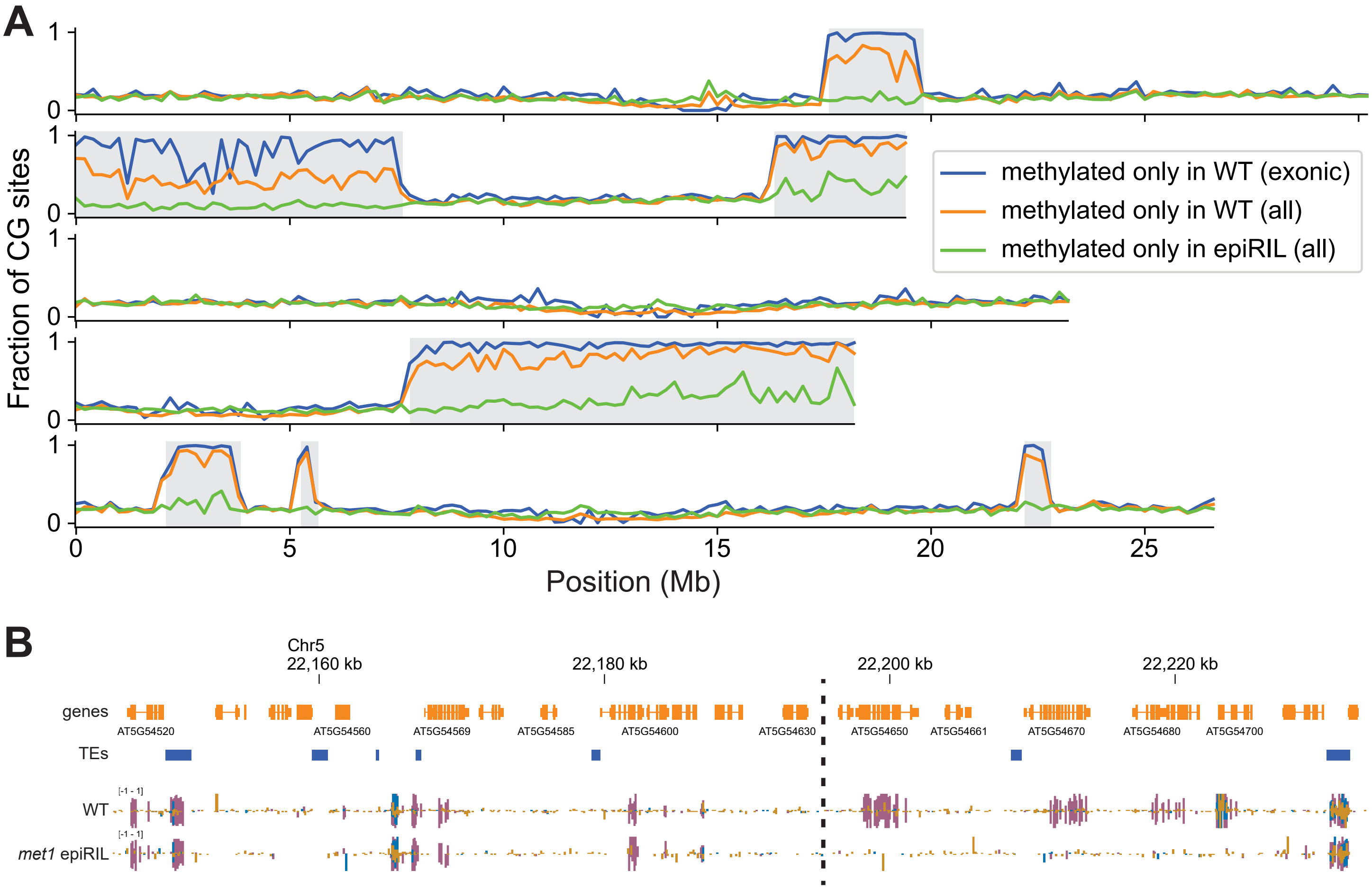

### Supplemental Figure S15. Defining *met1* epiRIL methylation loss regions

**A**, Fraction of CG sites which are methylated in WT but not in the *met1* epiRIL or vice versa. Seven large genomic windows were identified as losing almost all exonic mCG in the *met1* epiRIL (grey) and were used in Supplemental Figure S7F. **B**, Example genome browser view of the *met1* epiRIL breakpoint on Chr5. Gene body CG methylation is present in the *met1* epiRIL on the left side but not on the right. Purple=mCG, blue=mCHG, yellow=mCHH.
